# DOT1L stimulates MYC/Mondo transcription factor activity by promoting its degradation cycle on chromatin

**DOI:** 10.1101/2024.02.06.579191

**Authors:** Gian P. Sepulveda, Ekaterina S. Gushchanskaia, Alexandra Mora-Martin, Ruben Esse, Iana Nikorich, Ainhoa Ceballos, Julian Kwan, Benjamin C. Blum, Prakruti Dholiya, Andrew Emili, Valentina Perissi, Maria D. Cardamone, Alla Grishok

## Abstract

The proto-oncogene c-MYC is a key representative of the MYC transcription factor network regulating growth and metabolism. MML-1 (Myc- and Mondo-like) is its homolog in *C. elegans*. The functional and molecular cooperation between c-MYC and H3 lysine 79 methyltransferase DOT1L was demonstrated in several human cancer types, and we have earlier discovered the connection between *C. elegans* MML-1 and DOT-1.1. Here, we demonstrate the critical role of DOT1L/DOT-1.1 in regulating c-MYC/MML-1 target genes genome-wide by ensuring the removal of “spent” transcription factors from chromatin by the nuclear proteasome. Moreover, we uncover a previously unrecognized proteolytic activity of DOT1L, which may facilitate c-MYC turnover. This new mechanism of c-MYC regulation by DOT1L may lead to the development of new approaches for cancer treatment.

## INTRODUCTION

c-MYC is a transcription factor that regulates pathways essential to cell metabolism, proliferation, and animal development.^1^ It belongs to the basic helix-loop-helix leucine zipper (bHLHZ) family of transcription factors and forms a heterodimer with MAX to bind to the canonical E-box promoter sequence.^2^ c-MYC has been shown to regulate proliferation, apoptosis, ribosome biogenesis, nucleotide metabolism, and the epithelial-to-mesenchymal transition (EMT).^1,3–6^ c-MYC is a member of a larger group of MYC proteins, including MYCN and MYCL, preferentially acting as transcription activators.^7^ Notably, MAX forms heterodimers with both transcriptional activators and repressors, and there is an additional protein homologous to MAX, MLX (Max-like protein x),^8^ that binds some of the same repressor proteins as MAX and two additional bHLHZ activator proteins, MLXIP (MLX interacting protein) or Mondo A and MLXIPL (MLX interacting protein-like) or Mondo B, also known as ChREBP (Carbohydrate response element binding protein).^7^ Together, MYC proteins, MAX/MLX proteins, MLXIP (Mondo) proteins, and MYC/Mondo antagonists form the MYC transcription factor network.^7^ This network is simplified in the nematode *C. elegans* where there are two separate interacting pairs of MYC/MAX-like heterodimers: MML-1(MYC and Mondo-like)/MLX-2 and MDL-1/MLX-1.^9–11^ Here, we focus on a new common mechanism of regulation of MML-1 and c-MYC by a conserved histone H3 lysine 79 (H3K79) methyltransferase DOT1 (DOT-1.1 in *C. elegans* and DOT1L in mammals).

c-MYC was thought to regulate transcription solely via its Transactivation Domain (TAD) at the E-box-containing promoters of target genes.^12^ However, this dogma has been challenged over the years,^13–16^ revealing a more complex dynamic of c-MYC function at promoters. Our study of the sole MYC/Mondo family representative in *C. elegans*, MML-1, demonstrates that its binding to DNA is not sufficient to exert an effect on the transcription of target genes and that an additional regulatory step must be required.^17^ This work is consistent with an earlier study of transcription activation in response to a controlled increase in cellular c-MYC levels that did not show saturation at high c-MYC concentrations and suggested a model whereby c-MYC binding to promoters was not a rate-limiting step in gene regulation.^14^ This study proposed that c-MYC gets activated subsequently to promoter binding and that the activated form of c-MYC dissociates from promoters rapidly, likely by being degraded.^14^ Indeed, further research demonstrated that c-MYC degradation on chromatin by the nuclear proteasome is important for its proper function.^18–21^

While c-MYC is a well-recognized oncoprotein upregulated in almost 70% of cancers,^1^ the abnormal expression of DOT1L in solid tumors like breast, ovarian, prostate, and colorectal cancer has only been recently uncovered,^22–29^ although its role in MLL-rearranged leukemias has been studied for the past 20 years.^22,30–34^ Previous work has shown that DOT1L can cooperate with c-MYC and CBP/p300 in triple-negative breast cancer to activate the expression of genes needed for epithelial-mesenchymal transition (EMT) like *SNAI1*, *ZEB1,* and *ZEB2*.^28^ In neuroblastoma, MYCN and DOT1L were shown to interact and activate transcription of MYCN target genes.^35^ In *C. elegans*, the cooperation between DOT-1.1 and MML-1 has also been proposed due to their co-localization at select promoters and similar effects on hermaphrodite-specific neuron migration.^17^ There is also a strong correlation between H3K79me2 localization deposited by DOT1L and c-MYC binding to E-box promoters.^36^

While H3K79 methylation is found close to the transcription start sites (TSS) and within the bodies of actively transcribed genes,^37^ the causative and mechanistic link between H3K79 methylation and transcription is not fully defined.^38^ Even though H3K79 methylation is often associated with active gene transcription, a recent study showed that mutation of the DOT1L catalytic histone methyltransferase (HMT) domain only affected a subset of genes that were downregulated in a full knockout of DOT1L in embryonic stem cells, suggesting an additional HMT-independent mechanism of gene regulation by DOT1L during development.^39^ Additionally, DOT1L/DOT-1.1 was shown to regulate genes both positively^38,40,41^ and negatively^42–44^ and was reported to affect transcription initiation^45^ and elongation.^39^ It is clear that despite its wide pattern of expression, DOT1L’s molecular function is strongly influenced by the cell, tissue, and developmental context.

Here, we present evidence that metazoan DOT1 proteins, DOT-1.1 and DOT1L, strongly affect the MYC transcription factor network in *C. elegans* and mammals, respectively. Moreover, we show that DOT1L works with the ubiquitin-proteasome system (UPS) to regulate c-MYC occupancy on DNA, which is critical for c-MYC function in transcription regulation. Additionally, we uncover the novel ability of DOT1L to proteolytically cleave c-MYC which may aid with its subsequent degradation by the nuclear proteasome to maintain an active transcription cycle.

## RESULTS

c-MYC and DOT1L have recently been functionally connected in several cancer models.^23,26,28^ At the same time, we found links between *C. elegans* MML-1 and DOT-1.1.^17^ Here, we use both the nematode and mammalian cell culture systems to investigate this connection further.^10,19^

### ceDOT1L complex and MML-1 co-regulate genes

To determine whether MML-1, DOT-1.1, and ZFP-1, a DOT-1.1 binding partner and AF10 homolog,^42^ co-localize genome-wide, we performed MML-1 chromatin immunoprecipitation followed by deep sequencing (ChIP-seq) experiments in wild-type worms with an N-terminus-specific anti-MML-1 antibody **(Figure 1A)**.^17^ The overlap between MML-1 localization and the previously annotated ZFP-1/DOT-1.1 peaks^42^ was very significant **(Figure 1A)**, although the number of detectable MML-1 peaks was much lower compared to those of DOT-1.1 and ZFP-1 suggesting that DOT-1.1 and ZFP-1 have a larger effect on *C. elegans* genome.

**Figure 1.**
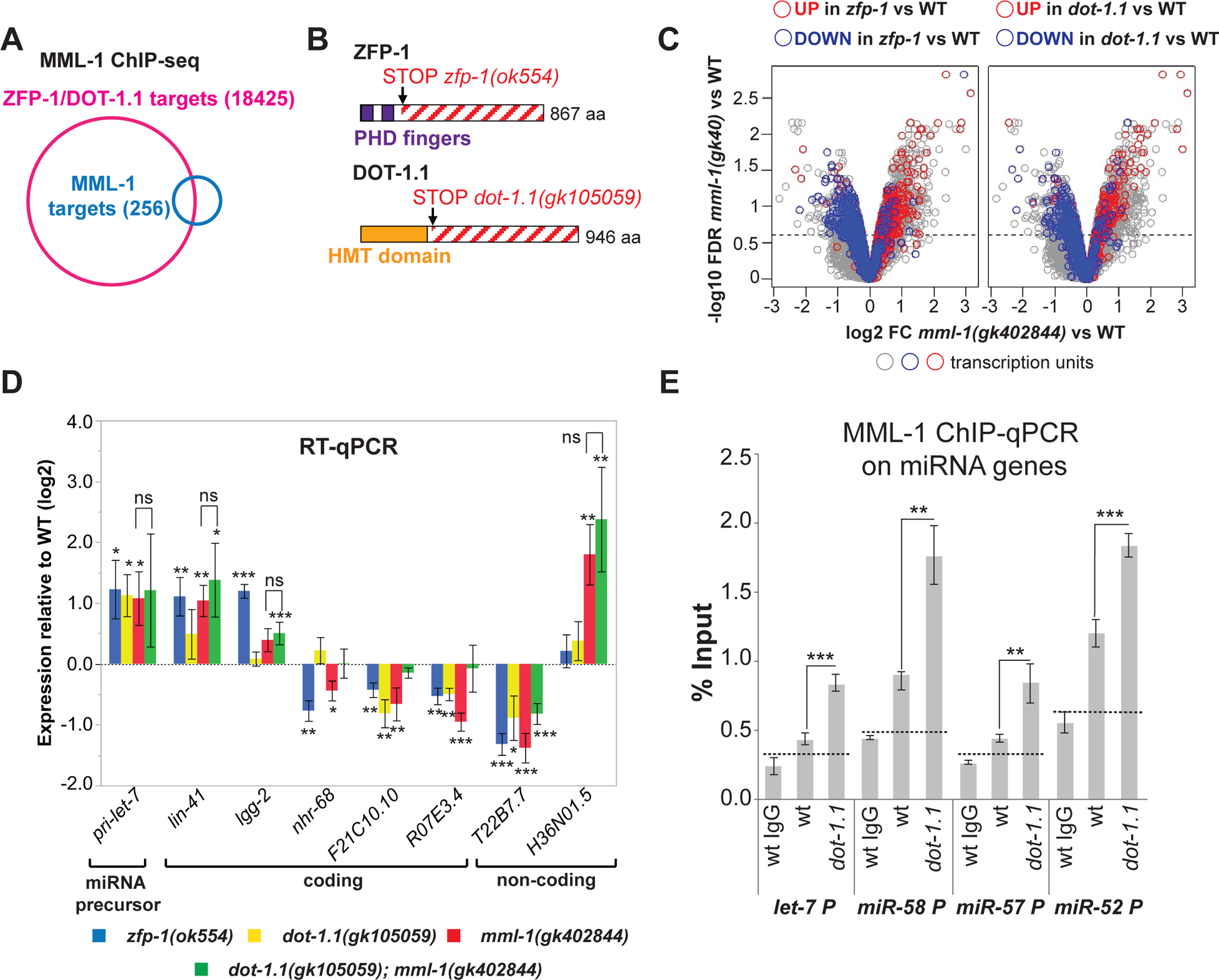
ceDOT1L complex and MML-1 co-regulate genes. **A.** Venn diagram showing an overlap between combined ZFP-1/DOT-1.1 ChIP-chip data^42^ and MML-1 ChIP-seq. P value less than 10^−6^ (Fisher’s exact test). **B.** *zfp-1* and *dot-1.1* mutant alleles used in microarray experiments in Figure 1C. Schematic of the effects of mutations leading to truncated proteins that do not interact with the respective (DOT-1.1 or ZFP-1) partner **C.** A volcano plot demonstrating changes in expression of ZFP-1/DOT-1.1-bound genes in *mml-1(gk402844)* compared to wild type (all open circles). Genes also changing expression in *zfp-1(ok554)* (left) or *dot-1.1(gk105059)* (right) are colored. **D.** RT-qPCR expression analyses of indicated coding and non-coding genes in single mutants and *dot-1.1; mml-1* double mutant strain compared to wild type. Specific genes were chosen based on microarray analyses shown in C, with the exception of *let-7*. H36N01.5 is an antisense transcript at the *daf-16* locus. Asterisks *, ** and *** denote P values (Student’s t-test) less than 0.05, 0.01 and 0.001, respectively, mutant *vs* wild type. Values are mean ± standard error obtained from 2 to 7 independent experiments. **E.** MML-1 ChIP-qPCR experiment showing MML-1 enrichment at indicated promoters (% Input) in wild type and mutant L3 larvae. Dashed lines denote ChIP background, based on ChIP with unrelated IgG. Error bars denote SD of triplicate qPCR runs. Asterisks *, ** and *** denote P values (Student’s t-test) less than 0.05, 0.01 and 0.001, respectively, mutant *vs* wild type or as indicated.

Next, we examined gene expression changes in *zfp-1(ok554), dot-1.1(gk105059)*, and *mml-1(gk402844)* mutant strains compared to wild-type. The *dot-1.1(gk105059)* allele harbors a stop codon after the methyltransferase domain of DOT1.1, producing a truncated protein that only contains this domain **(Figure 1B, bottom)**.^46^ This mutated DOT-1.1 protein lacks a ZFP-1-interacting domain, whereas *zfp-1(ok554)* produces a truncated protein missing a portion interacting with DOT-1.1 **(Figure 1B, top)**.^42,47^ We have previously shown that DOT-1.1 and ZFP-1 interact and colocalize at target genes and that mutation of ZFP-1 causes a reduction in H3K79me2.^42^ The *mml-1(gk402844)* mutant does not produce the MML-1 protein and therefore is considered null.^17^ A comparison of global gene expression changes in *zfp-1* and *dot-1.1* partial loss-of-function mutants (relative to wild-type) to the changes seen in *mml-1(gk402844)* clearly indicates co-regulation of genes by ZFP-1/DOT-1.1 and MML-1 **(Figure 1C)**, with similar sets of genes downregulated and upregulated in the *zfp-1*, *dot-1,* and *mml-1* mutants.

The co-regulation of genes co-occupied by ZFP-1/DOT-1.1 and MML-1 indicates that they either work independently, in parallel, or in the same pathway. To differentiate between these possibilities, we compared gene expression changes observed in single *mml-1(gk402844)* or *dot-1.1(gk105059)* mutants to those detected in the double mutant strain: *dot-1.1(gk105059); mml-1(gk402844)*. We found that changes in expression in the double mutant strain did not exceed changes in expression seen in the *mml-1(gk402844*) null mutant **(Figure 1D)**. This genetic relationship indicates that ZFP-1/DOT-1.1 and MML-1 act in the same pathway.

Similar to its mammalian homolog c-MYC, MML-1 either activated or repressed its direct targets in *C. elegans* larvae **(Figure 1D)**. For example, the null *mml-1* mutation led to an upregulation of the *let-7* miRNA precursor **(Figure 1D, left)** indicating that MML-1 acts as a repressor of *let-7* transcription. Since DOT-1.1 acts in the same pathway as MML-1, as expected, mutations in *dot-1.1* and *zfp-1* also led to *let-7* upregulation. Next, we considered the possibility that DOT-1.1 promotes MML-1 binding to target genes. For this, we evaluated MML-1 occupancy at its direct miRNA target promoters in the *dot-1.1* mutant background **(Figure 1E)**. Surprisingly, MML-1 showed increased promoter occupancy at the *let-7* promoter in *dot-1.1(gk105059)* **(Figure 1E)**. This increase in occupancy was also seen at other miRNA promoters. These results indicate that DOT-1.1 is not required for MML-1 interaction with DNA. Instead, in the *dot-1.1(gk105059)* background, there is an excessive accumulation of apparently inactive MML-1 at target promoters.

### DOT1L and c-MYC co-regulate genes in triple-negative breast cancer cells

To determine whether the mammalian homologs of DOT-1.1 and MML-1, DOT1L and c-MYC, respectively, co-regulate genes similarly to their *C. elegans* counterparts, we sought to create DOT1L knockout triple-negative breast cancer cells (MDA-MB-231) via CRISPR/Cas9.^48–50^ The MDA-MB-231 cell line was the first used to determine the interaction and cooperation between DOT1L and c-MYC in the regulation of the specific EMT genes *SNAI1*, *ZEB1,* and *ZEB2*.^28^

We used a single guide RNA (sgRNA) directed to the first exon of the *DOT1L* gene in our experiments **(Figure 2A)**. We confirmed the depletion of DOT1L protein in the sgRNA-treated cell population #2 via an IP-western blot **(Supplemental Figure 1A, last lane)**. It was not possible to generate a viable complete *DOT1L* knockout isolate from the sgRNA-treated cell populations in repeated attempts and with different sgRNAs. However, we were successful in propagating a viable *DOT1L* haploinsufficient MDA-MB-231-derived cell line. It contains an early stop codon in one *DOT1L* allele, and a 15-bp (5 amino acid) in-frame deletion in the other allele, as determined by deep sequencing of the DNA locus targeted by the sgRNA and the RNA-seq data **(Supplemental Figure 1B)**. We found that the histone methyltransferase activity of DOT1L is significantly reduced in the *DOT1L KO/+* cells **(Figure 2B)**. Moreover, these cells grow significantly slower than the original MDA-MB-231 cell line **(Figure 2C)**.

**Figure 2.**
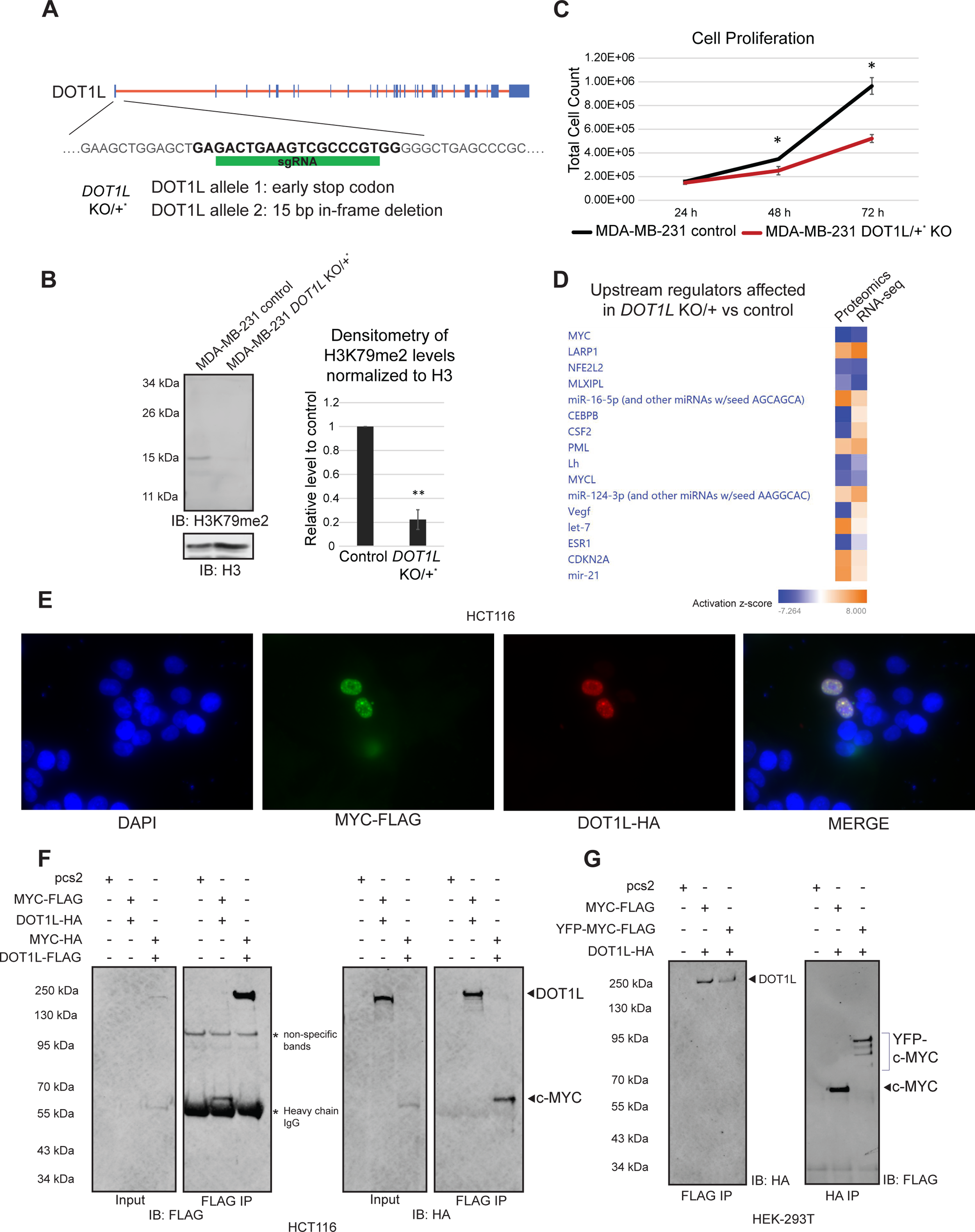
Mammalian DOT1L and c-MYC co-regulate genes in triple negative breast cancer cells. **A.** A schematic of the *DOT1L* locus and the sgRNA used to target the first exon of *DOT1L* for CRISPR/Cas9 gene editing. A clonal MDA-MB-231 *DOT1L KO/+* line was generated with one allele containing an early stop codon in the *DOT1L* locus, and the other allele containing a 15 base-pair in-frame deletion. **B.** *DOT1L KO/+* MDA-MB-231 cells have decreased H3K79me2 levels on western blot (N>5). **C.** Cell counting proliferation assays show that the *DOT1L KO/+* MDA-MB-231 cells grow significantly slower than the wild-type counterparts (N=2, with three technical replicates). **D.** Top upstream regulators affected in the *DOT1L KO/+* MDA-MB-231 cells compared to the wild-type counterpart as analyzed by Ingenuity Pathway Analysis. Proteomic analysis was performed on the *DOT1L* KO heterogenous population and compared to wild-type MDA-MB-231 (N=5). Bulk RNA-seq was done on the clonal *DOT1L KO/+* MDA-MB-231 cell line and compared to the wild-type controls (N=4). Blue squares represent a negative activation z-score indicating a downregulated pathway in the *DOT1L KO/+* MDA-MB-231 cells compared to wild-type, while orange squares represent a positive activation z-score indicating an upregulated pathway in the *DOT1L KO/+* MDA-MB-231 cells compared to wild-type. Proteomic and RNA-seq analyses for the top differentially regulated canonical pathways are mostly in agreement with one another. The MYC pathway is the top pathway downregulated in the differentially regulated canonical pathways in the *DOT1L KO/+* MDA-MB-231 cells compared to the wild-type controls as analyzed by Ingenuity Pathway Analysis. **E.** Overexpressed DOT1L and c-MYC can co-localize in distinct clusters in the nucleus in HCT116 cells. **F.** Overexpressed DOT1L immunoprecipitates with overexpressed c-MYC in HCT116 cells. Conversely, c-MYC immunoprecipitates with overexpressed DOT1L. **G.** Overexpressed DOT1L immunoprecipitates with overexpressed c-MYC or YFP-c-MYC in HEK-293T cells. Conversely, overexpressed c-MYC and YFP-c-MYC immunoprecipitate with overexpressed DOT1L.

To determine what pathways are perturbed when DOT1L function is reduced, we performed proteomics and total RNA-seq analyses on the *DOT1L KO/+* and wild-type cells. Principal component analysis shows a clear separation between wild-type and *DOT1L KO/+* samples **(Supplemental Figure 1C & 1D)**. Ingenuity Pathway Analysis and gene set enrichment analysis (GSEA) of both the proteomics and RNA-seq data sets revealed top pathways perturbed in the mutant **(Supplemental Figure 1E & 1F, Supplemental Table 1 & Supplemental Table 2).** The predominant effect was the downregulation of the cell cycle and growth-related processes in the *DOT1L KO/+* cells in both proteomic and RNA-seq datasets analyzed. Interestingly, EIF2 signaling was the most downregulated canonical pathway in the *DOT1L KO/+* cells indicating that mRNA translation was perturbed **(Supplemental Figure 1E)**. Furthermore, analyzing the most perturbed upstream regulator networks using Ingenuity Pathway Analysis software, we identified the c-MYC pathway as the top hit **(Figure 2D)**. GSEA of our RNA-seq data also shows that the hallmark c-MYC targets are downregulated in the *DOT1L KO/+* cells **(Supplemental Figure 1G & 1H).** Importantly, pathways governed by other members of the MYC transcription factor network like MLXIPL (Mondo B) and MYCL^51^ were also downregulated in both proteomic and RNA-seq analyses **(Figure 2D)**. Taken together, our genome-wide analyses indicate that the reduction of DOT1L function perturbs the c-MYC pathway and suggest that DOT1L and c-MYC co-regulate genes in triple-negative MDA-MB-231 breast cancer cells.

Earlier, it was shown that DOT1L and c-MYC co-immunoprecipitated in HEK-293 cells.^28^ To confirm this in other cell types, we looked at the colocalization of DOT1L and c-MYC in the human colorectal cancer cell line, HCT116, via overexpression of the two proteins **(Figure 2E)**. Interestingly, apart from their nuclear localization, DOT1L and c-MYC appeared to sometimes localize in bright, distinct clusters, providing further evidence of the two proteins co-regulating genes in a complex. Furthermore, overexpressed DOT1L co-immunoprecipitated with overexpressed c-MYC in HCT116 cells **(Figure 2F, lane 2)**. Conversely, overexpressed c-MYC co-immunoprecipitated with overexpressed DOT1L **(Figure 2F, lane 3)**. Similarly, in HEK-293T cells, overexpressed DOT1L co-immunoprecipitated with overexpressed c-MYC, as well as overexpressed YFP-c-MYC, confirming the previous study **(Figure 2G, left panel; Supplemental Figure 1I)**.^28^ Overexpressed c-MYC and YFP-c-MYC also co-immunoprecipitated with overexpressed DOT1L **(Figure 2G, right panel; Supplemental Figure 1I)**.

### DOT1L depletion leads to decreased c-MYC target gene expression but increases c-MYC occupancy at the target promoters

Next, we chose a panel of eight previously characterized c-MYC target genes from the RNA-seq data to confirm their reduced expression in *DOT1L KO/+* cells by RT-qPCR. The *miR-17* gene is a direct target of c-MYC activation and is essential in regulating E2F1 expression.^52^ Additionally, miR-17 inhibits *c-myc* mRNA expression in a negative feedback loop,^53^ and is essential in maintaining a neoplastic state crucial for cancer proliferation and survival.^54^ *PAICS* and *PPAT*, genes coding for two enzymes essential for de novo purine synthesis share the same bidirectional promoter and were shown to be directly upregulated by c-MYC.^55,56^ Moreover, PAICS inhibition was shown to suppress c-MYC-driven proliferation.^57^ c-MYC hallmark targets **(Supplemental Figure 1G & 1H)** *HK2*, *MAD2L*1, and *SMARCC1* are involved in glucose metabolism, cell cycle progression, and chromatin remodeling, respectively.^56,58–62^ Indeed, the expression of all eight chosen c-MYC targets was decreased in the *DOT1L KO/+* cells compared to the original cell line **(Figure 3A)**.

**Figure 3.**
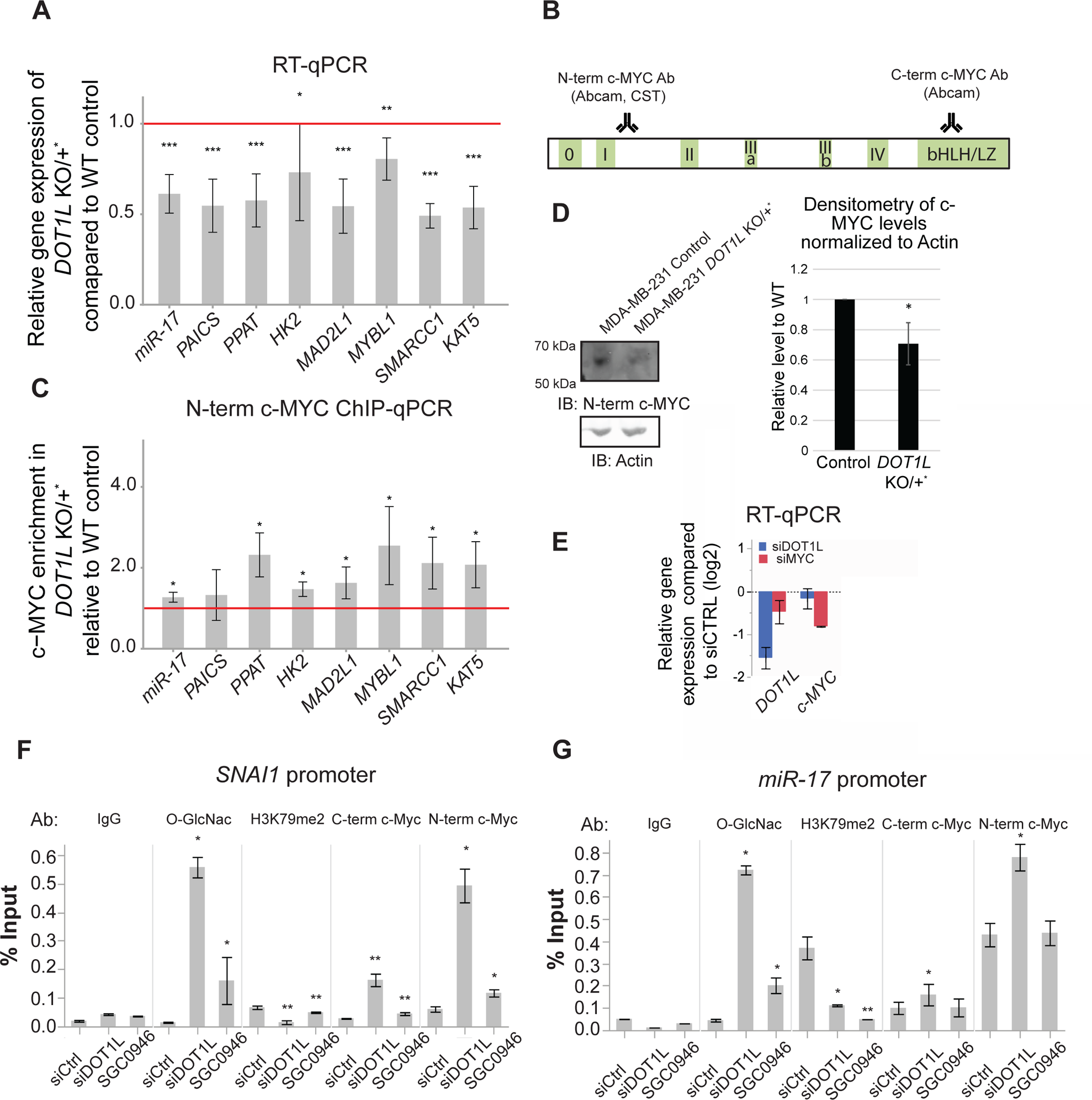
Depletion in DOT1L leads to decreased c-MYC target gene expression, but increased c-MYC occupancy. **A.** RT-qPCR of c-MYC target genes show a decrease in mRNA expression in *DOT1L KO/+* cells compared to wild-type cells (N= 8). All eight of the selected target genes have decreased mRNA expression. **B.** A schematic of the c-MYC protein and the corresponding regions of antibody recognition, **C.** c-MYC ChIP-qPCR of c-MYC target gene promoters shows an increase in c-MYC occupancy despite the decrease in gene expression. All eight target gene promoters had increased c-MYC occupancy (N=4). **D.** Western blot shows that the *DOT1L KO/+* cells have decreased c-MYC protein levels compared to the wild-type controls (N=3). Densitometry analysis of the c-MYC band is shown on the right. **E.** RT-qPCR shows that acute DOT1L and c-MYC knockdown leads to downregulation of their corresponding gene expression. **F & G.** c-MYC occupancy at the *SNAI1* and *miR-17* promoters is increased upon acute depletion of DOT1L via siRNA (N=3). The occupancy of *O*-GlcNAcylated proteins is also increased upon DOT1L knockdown. H3K79me2 in both promoters is decreased upon siDOT1L and upon treatment with a DOT1L histone methyltransferase (HMT) inhibitor, SGC0946. Inhibition of HMT activity did not increase c-MYC occupancy at the two target promoters to the same extent as DOT1L knockdown. Asterisks *, ** and *** denote P values (Student’s t-test) less than 0.05, 0.01 and 0.001, respectively.

To determine whether DOT1L and c-MYC co-regulate our chosen target genes in another cancer cell line we used HCT116 human colorectal carcinoma cells. We generated an auxin-inducible degron (AID)^63–65^ system targeting c-MYC in these cells. 50 minutes after auxin administration, there was near total depletion of the c-MYC-AID protein **(Supplemental Figure 2A)**. To see the effect of c-MYC depletion on its target gene expression, we treated cells with auxin for 4 hours before performing RT-qPCR. There was a trend for reduction of gene expression in most c-MYC target genes tested in MDA-MB-231 *DOT1L KO/+* cells, further confirming that these genes are activated by c-MYC **(Supplemental Figure 2B)**. Next, we performed DOT1L knock-down in HCT116 cells using shRNA and observed a decrease in expression of the tested c-MYC target genes as well **(Supplemental Figure 2C)**. DOT1L knockdown via shRNA led to a decrease in H3K79me2 indicating that shRNA treatment was effective **(Supplemental Figure 2D)**. These results indicate that cooperation between c-MYC and DOT1L in gene regulation is not limited to breast cancer.

We then used c-MYC ChIP coupled with qPCR using the N-terminal c-MYC antibody **(Figure 3B)** to look at c-MYC occupancy at the promoters of the tested target genes in MDA-MB-231 *DOT1L KO/+* cells compared to the control cells. Surprisingly, we found that c-MYC occupancy was increased in the *DOT1L KO/+* cells **(Figure 3C; Supplemental Figure 2E & 2F)**. This result is similar to that we observed for MML-1 in the *dot-1.1* mutant background in *C. elegans* **(Figure 1E)**. Notably, the increase in c-MYC occupancy occurs despite the slight decrease of c-MYC protein (and mRNA) levels in MDA-MB-231 *DOT1L KO/+* cells compared to the control cells **(Figure 3D)**.

It has been previously proposed that c-MYC binding to promoters is not sufficient for target gene regulation and that a c-MYC activating event is needed to make it competent in affecting transcription.^14,20,66,67^ Moreover, it was suggested that c-MYC binding to promoters is not static and that the on-off rate of c-MYC binding might be important for its effect on RNA Pol II-mediated transcription.^14,67^ The increase of inactive c-MYC occupancy at its target promoters in *DOT1L KO/+* cells indicates that DOT1L plays a role in the activation of c-MYC after its binding to DNA and in its cycling at chromatin.

Importantly, acute depletion of DOT1L in MDA-MB-231 via siRNA **(Figure 3E)** also leads to an increase in c-MYC occupancy at the *SNAI1* and *miR-17* gene promoters **(Figure 3F & 3G)**, as determined by ChIP-qPCR with two antibodies against c-MYC **(Figure 3B)**. c-MYC is modified by an *O*-linked N-acetylglucosamine (*O*-GlcNAc)^68^ and we used ChIP-qPCR with an antibody specifically recognizing *O*-GlcNAc as another readout of elevated c-MYC occupancy upon DOT1L siRNA treatment **(Figure 3F & 3G)**.

We also performed c-MYC and *O*-GlcNAc ChIP-qPCR experiments using cells treated with a specific DOT1L methyltransferase inhibitor SGC0946^69^ and determined that the increase in c-MYC occupancy is partially independent of the HMT activity of DOT1L since inhibitor treatment^71^ did not lead to the robust increase in c-MYC promoter occupancy compared to the effect of siDOT1L depletion **(Figure 3F & 3G)**. Importantly, SGC0946 treatment caused a significant reduction in H3K79me2 levels at the studied promoters **(Figure 3F & 3G),** as expected. Also, inhibitor treatment did not perturb DOT1L localization to the nucleus (**Supplemental Figure 3A-3C)**.

Overall, our results obtained using cancer cells indicate that under conditions of DOT1L depletion, c-MYC has increased occupancy at target promoters, but lacks the ability to activate gene transcription.

### DOT1L works with the ubiquitin-proteasome system to activate c-MYC-driven gene transcription

Importantly, previous studies have implicated the proteasome in promoting the transcription of c-MYC target genes.^18,20,70,71^ It was found that the timely degradation of c-MYC at target promoters was essential for the c-MYC-dependent increase in target gene transcription.^18^ Furthermore, this regulation of transcription factors by the nuclear proteasome is not limited to c-MYC.^72,73^

*C. elegans* MML-1 is likely regulated by a similar mechanism, based on our reported observations. ^17^ Specifically, while investigating a strong loss-of-function *mml-1(ok849)* deletion mutant that exhibited a reduced lifespan^74^ with available anti-MML-1 antibody,^10^ surprisingly, we detected expression of the mutant protein. ^17^ The *mml-1(ok849)* deletion allele produced an internally truncated protein where the N-terminal and C-terminal portions of the protein were fused in-frame **(Supplemental Figure 4C)**. Since the C-terminus retained the bHLHZ domain, MML-1(ok849) binding to target promoters was not compromised. ^17^ In fact, the nonfunctional mutant protein exhibited increased promoter occupancy compared to wild-type MML-1, ^17^ similar to the accumulation of inactive MML-1 observed in *dot-1.1* mutant worms **(Figure 1E)**.^19^ The deleted region in this mutant is therefore required for MML-1 activation following its DNA binding.^17^ Moreover, the inactive MML-1 protein with internal deletion appears to be more stable compared to the wild-type based on the immunoblots.^17^ This suggests that regulation of MML-1 stability and turnover is important for its role in transcription and that DOT-1.1 may have a role in regulating MML-1 dynamics.

In a search for a possible role of DOT1L in protein turnover regulation, we conducted GSEA on a group of 1357 proteins that co-immunoprecipitated with DOT1L in HEK-293T cells in a previous study.^33^ We found an overrepresentation of the proteasome pathway, among others **(Supplemental Figure 4A)**. To determine whether DOT1L affects overall c-MYC protein stability, we used cycloheximide to inhibit translation. Surprisingly, we observed only a slight but insignificant decrease in total overall c-MYC stability and half-life in *DOT1L KO/+* cells **(Supplemental Figure 4B)**. Since we observe increased occupancy of c-MYC at the promoters in *DOT1L KO/+* **(Figure 3C)** and upon *DOT1L* siRNA **(Figure 3F & 3G)**, the effect of DOT1L depletion on c-MYC abundance appears to be localized to chromatin.

To determine whether DOT1L activates c-MYC target genes by facilitating its timely degradation via the proteasome, we treated both the control and *DOT1L KO/+* MDA-MB-231 cells with the proteasome inhibitor MG132 **(Figure 4A)**. Treatment with MG132 led to an elevation in global protein ubiquitination and an increase in full-length and ubiquitinated c-MYC protein levels, as expected **(Figure 4B)**. Despite the increase in c-MYC protein levels **(Figure 4B, left blot)**, most c-MYC target genes that were dependent on DOT1L for their full activation **(Figure 4C & 3A)** showed decreased expression in MG132-treated MDA-MB-231 cells **(Figure 4D)**, confirming the importance of c-MYC turnover in regulating target gene expression.

**Figure 4.**
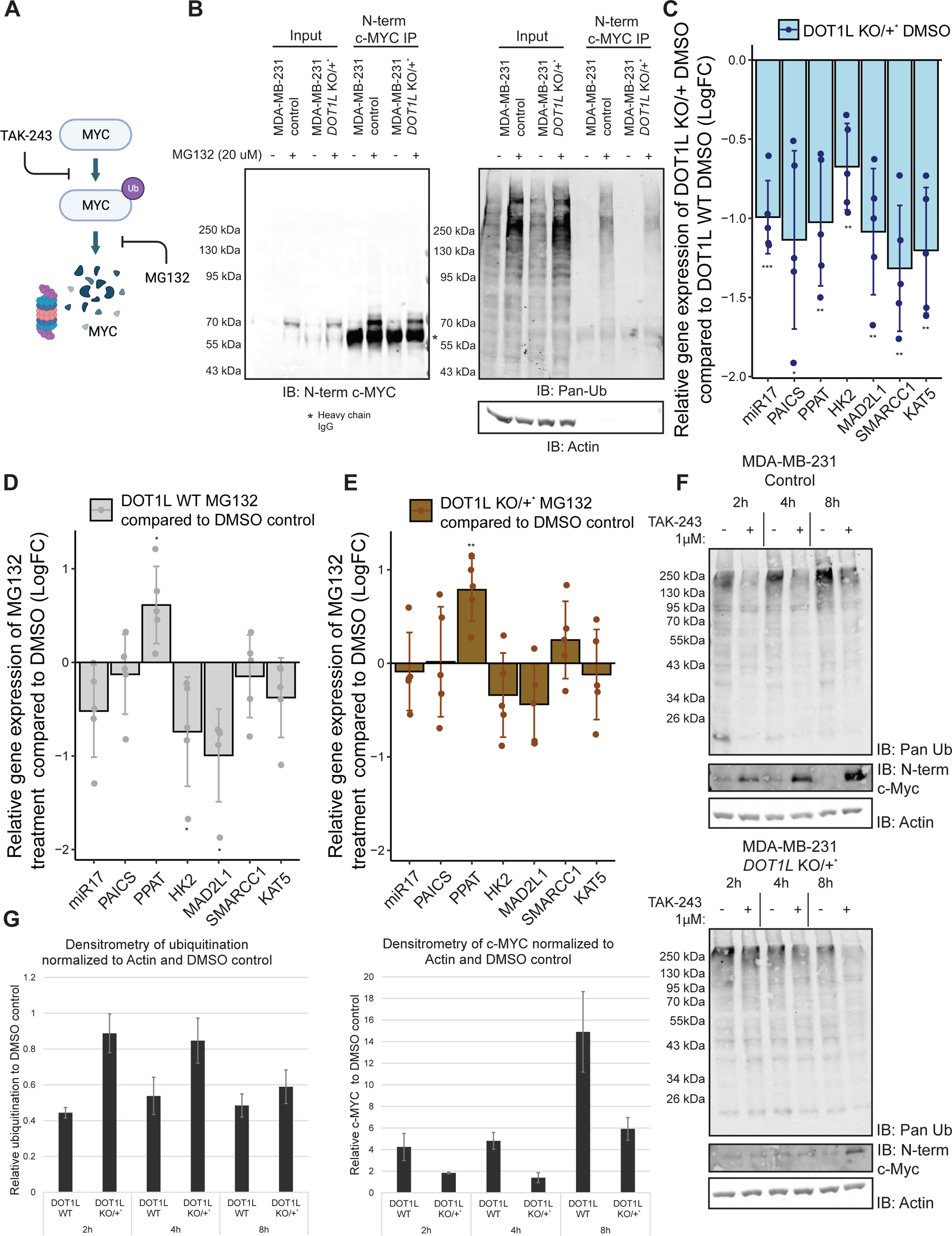
DOT1L works with the ubiquitin-proteasome system to activate c-MYC driven gene transcription. **A.** A schematic of the degradation process of c-MYC. This schematic was created with BioRender.com. **B.** Treatment of wild-type and *DOT1L KO/+* cells with 20 μM of MG132 leads to an increase in total ubiquitinated proteins, as well as c-MYC protein levels (full length and ubiquitinated) after 4 hours of treatment (N=5). **C.** Consistent with non-DMSO treated cells, c-MYC target gene expression is decreased in the *DOT1L KO/+* cells compared to the wild-type control cells under 4 hours of DMSO **D & E.** 4-hour treatment with 20 μM MG132 leads to a decrease in c-MYC target gene expression in the control cells (N=5). *DOT1L KO/+* cells have a dampened decrease in c-MYC target gene expression after MG132 treatment, indicating that the proteasome and DOT1L are in the same pathway for c-MYC activation. **F.** TAK-243 chase experiments show that proteasomal clearing of ubiquitinated substrates is delayed in *DOT1L KO/+* cells. c-MYC stabilization after ubiquitination inhibition is also delayed in the mutant cells compared to wild-type. **E.** Densitometry of c-MYC bands and ubiquitinated smears in **F** normalized to Actin and the respective DMSO controls.

Compared to the unmodified MDA-MB-231 cells, in the *DOT1L KO/+* background, MG132 treatment had less effect on gene expression **(Figure 4E)**. Thus, genes sensitive to proteasome inhibition in control cells lost this sensitivity upon partial DOT1L knockout. Note that these genes were already downregulated in *DOT1L KO/+* compared to the unmodified cells **(Figure 4C)**. This indicates that DOT1L and the proteasome work in the same pathway and that further inhibition of the proteasome in *DOT1L KO/+* cells does not lead to a further decrease in gene expression.

To determine whether the functionally important region deleted in *C. elegans* MML-1(ok849) was relevant for regulation via the proteasome, we used CRISPR/Cas9 to tag the endogenous *mml-1* gene with the FLAG tag **(Supplemental Figure 4C)**. We introduced the tag into the wild-type strain as well as into the *mml-1(ok849)* mutant coding for a more stable inactive protein with internal deletion.^17^ Next, we combined both tagged strains with the mutation in the proteasomal ubiquitin receptor gene *rpn-10*. The *rpn-10* loss-of-function mutants are viable but defective in proteasome function and exhibit an accumulation of ubiquitinated proteins targeted for degradation.^75,76^ Notably, we observed smearing above the MML-1::FLAG band detected by immunoblotting in the proteasomal mutant *rpn-10(ok1865),* consistent with the accumulation of ubiquitinated but not degraded MML-1::FLAG isoforms **(Supplemental Figure 4D, blue bracket)**.^19^ The endogenously tagged MML-1(ok846)::FLAG protein was detected as well **(Supplemental Figure 4D)**. However, in the *rpn-10(ok1865)* mutant background we did not observe a smear above the MML-1(ok846)::FLAG band **(Supplemental Figure 4D, red arrow)**. This indicates that MML-1(ok846) is less prone to ubiquitin-mediated proteasomal degradation. Moreover, since MML-1(ok846) is inactive,^17,74^ there is a correlation between MML-1 activity and its degradation by the proteasome, just like in the case of c-MYC in the triple-negative breast cancer system.

To determine whether DOT1L depletion influences the dynamics of ubiquitinated protein turnover, we treated control and *DOT1L KO/+* MDA-MB-231 cells with TAK-243 **(Figure 4A)**, a small molecule inhibitor of the ubiquitin-activating enzyme (UAE), which is the main E1 enzyme of the ubiquitin conjugation pathway.^77^ Inhibition of general protein ubiquitination led to a decrease in ubiquitinated proteins in the control MDA-MB-231 cells, as early as 2 hours into treatment **(Figure 4F, upper panel)**. Accordingly, the protein levels of c-MYC also increased upon treatment, since the protein is not being ubiquitinated and subsequently degraded **(Figure 4F, upper panel)**. In contrast, DOT1L depletion led to a delayed response to TAK-243 treatment, with a decrease in the ubiquitinated protein smear only being apparent at 8h post-treatment **(Figure 4F, lower panel)**. The protein level increase of c-MYC was also delayed, only showing an appreciable increase at 8 hours as well **(Figure 4F, lower panel)**. The observed changes were quantified via densitometry **(Figure 4G)**. These results indicate that the general turnover of ubiquitinated proteins is perturbed in DOT1L depleted conditions.

Taken together, our data indicate that DOT1L helps facilitate c-MYC turnover via the ubiquitin-proteasome system. This turnover is essential in activating c-MYC-driven transcription of target genes. Moreover, we have evidence that DOT1L has a more general effect on protein turnover which slows down when DOT1L levels are reduced.

### DOT1L cooperates with Valosin-containing protein to activate c-MYC-driven gene transcription

A recent study of proteasomal inhibition effects on c-MYC revealed that MG-132 treatment can lead to the multimerization of c-MYC and its redistribution from active promoters to stalled replication forks.^78^ We were able to recapitulate c-MYC multimer formation upon MG-132 treatment in U2OS and HCT116 cells **(Supplemental Figure 4E)**. To determine whether DOT1L depletion alone leads to multimer formation, we looked at c-MYC localization in control MDA-MB-231 cells and *DOT1L KO/+* cells via immunofluorescence. Neither the wild-type nor the *DOT1L KO/+* cells had noticeable multimer formation under DMSO control conditions **(Supplemental Figure 4F, DMSO row)**. However, upon treatment with MG-132, both cell types had noticeable multimer formation, albeit smaller than the U2OS MG-132 treated cells **(Supplemental Figure 4F, MG-132 row)**. Therefore, MG-132 treatment and genetic DOT1L depletion have both common (reduced c-MYC target gene expression) and distinct (multimer formation) effects on c-MYC.

Next, we wanted to further narrow down the step in which DOT1L acts in the degradation cycle of c-MYC on chromatin. Interestingly, inhibition of the proteasomal degradation system via Valosin-containing protein (VCP) inhibitor NMS-873 **(Figure 5A)** does not lead to multimer formation.^78^ VCP is an AAA+ ATPase also known as p97 and Cdc48.^79,80^ VCP is an important part of the UPS responsible for the extraction of target proteins from the membranes^79,81^ and chromatin.^82,83^

**Figure 5.**
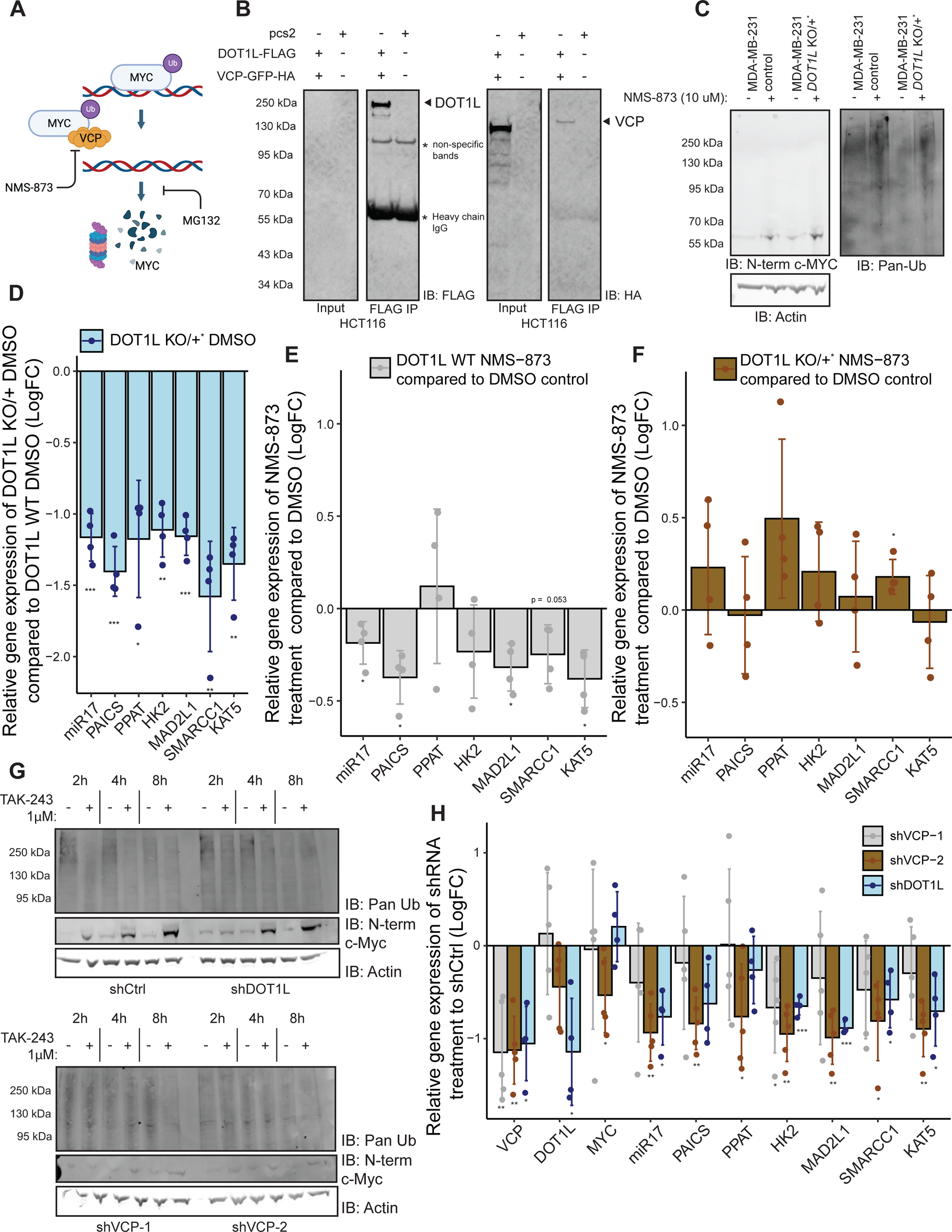
DOT1L cooperates with VCP to regulate c-MYC-driven gene transcription. **A.** A schematic of the proposed mechanism by which VCP extracts c-MYC from chromatin for degradation. This schematic was created with BioRender.com. **B.** Overexpressed VCP-GFP immunoprecipitates with overexpressed DOT1L in HCT116 cells (N=2). **C.** 6-hour treatment with 10 μM of NMS-873 leads to an increase in ubiquitinated proteins and c-MYC protein levels in both the wild-type control and *DOT1L KO/+* cells (N=4). **D.** Consistent with non-DMSO treated cells, c-MYC target gene expression is decreased in the *DOT1L KO/+* cells compared to the wild-type control cells under 6 hours of DMSO. **E & F.** 6-hour treatment with 10 μM NMS-873 leads to a significant decrease in c-MYC target gene expression in the control cells (N=4). This decrease is absent in *DOT1L KO/+* cells, indicating that VCP and DOT1L are in the same pathway for c-MYC activation. **G.** TAK-243 chase experiments show that proteasomal clearing of ubiquitinated substrates is delayed in MDA-MB-231 cells with shDOT1L and shVCP compared to shCtrl. c-MYC stabilization after ubiquitination inhibition is also delayed in the shDOT1l and shVCP cells compared to shCtrl. **H.** shRNA against VCP and DOT1L leads to a decrease in c-MYC target gene expression.

To determine whether DOT1L and VCP are present in the same complex, we performed immunoprecipitation experiments with overexpressed DOT1L and VCP. We determined that VCP-GFP was co-immunoprecipitated when DOT1L is pulled down in both HCT116 **(Figure 5B)** and HEK-293T cells **(Supplemental Figure 5A)**. Furthermore, VCP-GFP localized to both the cytoplasm and nucleus in HCT116 cells, suggesting that DOT1L, MYC and VCP may interact in the nucleus **(Supplemental Figure 5B)**.

Critically, VCP has been implicated in the activation of c-MYC target genes.^71^ It is thought that VCP is needed to extract ubiquitin-modified c-MYC from chromatin before its subsequent degradation and that this step is important for c-MYC target gene expression.^71^ To determine whether DOT1L acts in this step of c-MYC activation, we treated both the control and *DOT1L KO/+* MDA-MB-231 cells with the VCP inhibitor NMS-873. Our RNA-seq analysis showed no difference in VCP mRNA expression between *DOT1L KO/+* MDA-MB-231 cells and the control cells in untreated conditions **(Supplemental Table 1)**. Furthermore, there was no difference in the total VCP protein levels **(Supplemental Figure 5C)** or VCP hexamerization **(Supplemental Figure 5D)** between the control and *DOT1L KO/+* cells. Similar to our results from previous experiments, *DOT1L KO/+* cells had lower c-MYC target gene expression compared to control cells when under DMSO treatment conditions **(Figure 5D)**. Also, similar to proteasomal inhibition using MG132 **(Figure 4D)**, treatment with NMS-873 led to an increase in c-MYC protein levels and an increase in ubiquitinated proteins in both control and *DOT1L KO/+* cells **(Figure 5C)**. Despite the increase in c-MYC protein levels, VCP inhibition led to a decrease in c-MYC target gene expression in the control cells **(Figure 5E)**, consistent with published work.^71^ This decrease was absent in the *DOT1L KO/+* MDA-MB-231 cells **(Figure 5F)** suggesting that DOT1L and VCP act in the same pathway for c-MYC target gene activation.

To further confirm that DOT1L and VCP work in the same pathway, we used a short hairpin RNA approach. Knockdown of VCP, using shVCP-1 led to an increase in ubiquitinated proteins in both the control and *DOT1L KO/+* cells, similar to NMS-873 treatment **(Supplemental Figure 5E)**. shVCP-2 led to an increase in ubiquitinated proteins in the control cells but not the *DOT1L KO/+* cells **(Supplemental Figure 5E)**, consistent with VCP downregulation failure by shVCP-2 in the mutant cells **(Supplemental Figure 5F)**. Additionally, shDOT1L led to a decrease in H3K79me2 in MDA-MB-231 cells **(Supplemental Figure 5E)**.

To determine whether VCP depletion influences the dynamics of ubiquitinated protein turnover, similar to DOT1L depletion **(Figure 4F)**, we treated MDA-MB-231 cells harboring the shRNA against VCP with TAK-243. Inhibition of general protein ubiquitination led to a decrease in ubiquitinated proteins in the shControl cells, as early as 2 hours into treatment **(Figure 5G, upper panel).** shDOT1L treatment led to a delay in the response to TAK-243 treatment **(Figure 5G, upper panel)**, similar to the *DOT1L KO/+* cells **(Figure 4F, Supplemental Figure 5G)**. The shVCP treatment of the control MDA-MB-231 cells also resulted in a delay in the response to TAK-243, similar to the DOT1L depletion conditions **(Figure 5G, lower panel)**. shVCP treatment of the *DOT1L KO/+* cells did not delay the response to TAK-243 any further **(Supplemental Figure 5G)**, indicating that VCP and DOT1L work together in facilitating the turnover of ubiquitinated proteins. Furthermore, shVCP treatment led to a decrease in c-MYC target gene expression **(Figure 5H)**, similar to DOT1L depletion. Thus, our data show that DOT1L cooperates with VCP to facilitate the turnover of c-MYC, which activates gene expression.

### Proteolytic cleavage of MML-1 and c-MYC

We noted that, in *C. elegans*, the available multicopy transgenic MML-1::GFP::FLAG construct hardly produced a full-length protein in wild-type background. Rather, a ∼60 kDa C-terminal fragment appeared as the dominant form of the transgene-encoded protein **(Figure 6A)**. Importantly, in the *zfp-1(ok554)* mutant background with reduced DOT-1.1 localization to chromatin,^42^ we observed elevated expression of the full-length MML-1::GFP::FLAG with a concomitant reduction in the shorter C-terminal fragment levels **(Figure 6A)**. These changes are consistent with MML-1 cleavage and demonstrate that the DOT-1.1 complex has a role in this process. The size of the band indicates that the cleavage occurs somewhere in the proline-rich region of MML-1, which is missing in the inactive MML-1(ok846) that accumulates at the target promoters.^17^ This suggests that cleavage in the proline-rich region is important in the activation of MML-1.^19^

**Figure 6.**
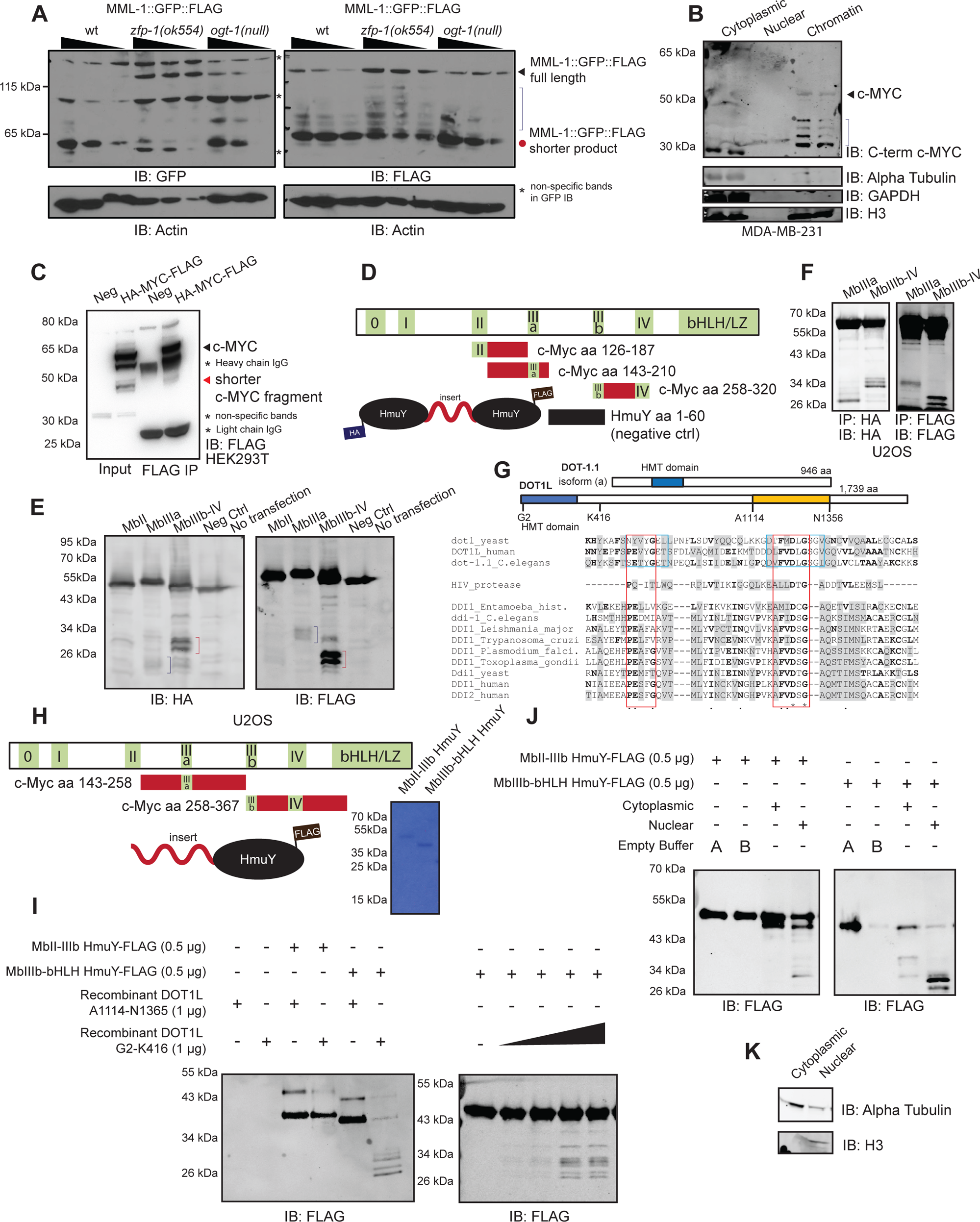
DOT1L has proteolytic activity and shorter c-MYC fragments observed in cells are the result of proteolytic cleavage. **A.** Transgenic MML::FLAG::GFP in wild-type worms produces a shorter MML::FLAG::GFP product that is stabilized to the full-length transgenic protein in the *zfp(ok544)* mutant background (N=2). This transgene is not stabilized in the *ogt-1(ok430)III* null worm. **B.** Cellular fractionation of MDA-MB-231 cells shows shorter c-MYC fragments in the chromatin fraction, detected by the C-terminal c-MYC antibodies (N=3). **C.** A consistent 45 kDa c-MYC fragment is produced upon transfection of c-MYC plasmids in HEK-293T cells, which can be pulled down via immunoprecipitation (N=3). **D.** A schematic of the protease-resistant HmuY constructs used to probe cleavage of different c-MYC fragments. The HmuY fragment aa 1-60 was used as a negative control for cleavage. **E.** Transfection of the HmuY constructs shows cleavage of c-MYC from aa 143-210, and aa 258-320, as detected by both the HA and FLAG antibodies (N=3). **F.** Immunoprecipitation of the N-terminal and C-terminal cleavage fragments of c-MYC produced from aa 143-210 and aa 258-320 using the HA and FLAG antibodies (N=2). **G.** A schematic of DOT-1.1 (*C. elegans*) and DOT1L (mammalian) proteins. A part of the HMT domain (aa 126-192 in human DOT1) of DOT1 family proteins aligns with the catalytic domain of the DDI family of aspartic proteases, which are related to HIV protease. Note the conservation of the catalytic aspartic acid (**D**S/T**G**) in DDI/HIV proteases and the **D**L**G** sequence in DOT1 enzymes. Blue boxes show regions of DOT1L that bind methyl donor S-Adenosyl methionine. Red boxes show the most homologous amino acid residues between DOT1 and DDI protein families. Bold letters indicate identical residues in at least one pair of DOT1 and protease protein family proteins. Grey shade indicates similar amino acid residues. CLUSTAL Omega tool was used to generate the multiple sequence alignment. **H.** A schematic of the protease-resistant HmuY constructs used for in vitro cleavage assays. Coomassie blue staining shows one predominant band for each in vitro construct. **I.** Cleavage by the N-terminus, but not C-terminus, of DOT1L occurs between aa 258-367 of c-MYC in vitro (N=3). Commercially available N-terminal DOT1L or purified C-terminal DOT1L expressed in E. coli was incubated with the respective HmuY constructs for 24 hours at 37 °C. **J.** The region between aa 143-258 of c-MYC is cleaved by the nuclear fraction of U2OS cells. The region between aa 258-367 of c-MYC is cleaved by both the cytoplasmic and nuclear fractions of U2OS cells to produce distinct cytoplasmic and nuclear cleavage bands. The respective HmuY constructs were incubated with 2.5 μg of U2OS cell fractions for 24 hours at 37 °C. **K.** Western blotting of alpha tubulin and H3 show that the cellular fractionation protocol successfully separated the cytoplasmic and nuclear fractions of U2OS cells.

There are precedents of protease-mediated cleavage of transcription factors leading to their activation^84–86^ and/or preceding their degradation by the proteasome.^87–90^ Interestingly, the HIV protease-related aspartic DNA-damage inducible (DDI) proteases have been implicated in the cleavage of NRF transcription factors: SKN-1 in *C. elegans*^84^ and NRF1(NFE2L1) in mammals.^91,92^ Importantly, NRF1 ubiquitination greatly facilitates cleavage by DDI2.^92,93^

To determine whether c-MYC is cleaved in vivo and in which cellular compartment this cleavage occurs, we performed cellular fractionation and observed smaller C-terminal c-MYC fragments in the chromatin fraction **(Figure 6B)**. Next, we used CRISPR/Cas9 to tag the endogenous c-MYC protein with a FLAG tag at the C-terminus in HCT116 colorectal cancer cells and observed that the endogenously tagged c-MYC protein also produces smaller c-MYC fragments that were stabilized when treated with a proteasome inhibitor **(Supplemental Figure 6A)**. Furthermore, overexpression of the exogenous HA- and FLAG-tagged c-MYC via transfections in HEK-293T cells consistently shows a ∼45 kDa C-terminal fragment that can be pulled down using a FLAG antibody **(Figure 6C)**. This C-terminal fragment can also be detected during exogenous c-MYC overexpression in HCT116 and U2OS cells **(Supplemental Figure 6B)**. We determined that this fragment is not a result of alternative translation from an internal start codon like in the case of s-MYC^15,94,95^ since mutation of the internal methionine amino acids still produces the ∼45 kDa band **(Supplemental Figure 6C)**. These findings suggest that the shorter c-MYC fragments are the result of proteolytic cleavage of the full-length protein.

We hypothesized that some c-MYC cleavage fragments could result from the cleavage around the middle portion of the protein. To facilitate cleavage detection, we developed a reporter construct using the *Porphyromonas gingivalis* HmuY protein as a scaffold due to its protease-resistant nature.^96,97^ Since this protein has been used in CleavEx^98^ in vitro technology, but not yet in cells, we first tested its expression in the HCT116 and U2OS cell lines. The protease-resistant HmuY protein was expressed strongly in HCT116 cells **(Supplemental Figure 6D)**. Next, we created reporter constructs using two tagged HmuY proteins flanking different c-MYC fragments spanning from MbII through MbIV, as well as a negative control insert containing the first 60 amino acids of the protease-resistant HmuY **(Figure 6D)**. These reporter constructs contained a nuclear localization signal. The construct comprising MbIIIb-MbIV showed a strong cleavage band when overexpressed in U2OS cells **(Figure 6E, red brackets)**. Notably, c-MYC cleavage by calpains at K298 located between MbIIIb and MbIV has previously been suggested, although specific enzymes responsible for it have not been identified.^99,100^ Our reporter results are consistent with calpain cleavage but do not exclude other possibilities. Interestingly, the construct spanning the region around MbIIIa also produced short cleavage fragments detected by both the N-terminus (HA) and C-terminus (FLAG)-specific antibodies **(Figure 6E, blue brackets)**. These cleavage products are further stabilized when the transfected cells are treated with MG132 **(Supplemental Figure 6E)**. Furthermore, the N-terminal and C-terminal fragments produced from the cleavage in the region spanning MbIIIa, as well as the fragments produced from the region spanning MbIIIb and MbIV, were pulled down via immunoprecipitation using the HA and FLAG antibodies **(Figure 6F)**.

We also constructed a reporter containing a FLAG-tagged Nanoluciferase at the N-terminus, a 117 amino acid spacer corresponding to residues 141 through 258 of the human c-MYC, which contains part of conserved Myc Box II (MbII) and the entire MbIIIa, and a YFP sequence at the C-terminus **(Supplemental Figure 6F)**. We introduced this reporter into HEK-293T cells and evaluated potential cleavage with FLAG-specific and GFP-specific immunoblotting. We detected N-terminal fragments using the FLAG antibody at around 27.5 kDa and 20 kDa, and detected a C-terminal fragment using the GFP antibody at around 35 kDa **(Supplemental Figure 6G)**. These fragments indicate that cleavage may occur at more than one region in the middle of c-MYC, reminiscent of the multiple c-MYC bands found in the chromatin fraction **(Figure 6B)**.

Taken together, our results indicate that both c-MYC and MML-1 are subject to a novel protease cleavage likely occurring at chromatin. Moreover, the MML-1 cleavage is promoted by the DOT-1.1 complex.

### DOT1L exhibits cleavage of c-MYC in vitro

The reduction in ubiquitinated protein turnover observed in DOT1L deficient cells **(Figure 4F)** is reminiscent of the similar effect of *DDI2* KO cells.^92^ This suggests that DOT1L and a DDI family protease may act in the same pathway and/or that DOT1L itself may act as a protease.

We performed multiple sequence alignment of the protease domains of DDI and HIV proteases and the Dot1 family proteins and, surprisingly, found conservation of the catalytic aspartic acid residue of the aspartic proteases in DOT1 proteins **(Figure 6G)**. The aspartic proteases of the HIV family exhibit the **D** (S/T) **G** consensus sequence at the catalytic site.^101^ This sequence aligned with the conserved **D**L**G** residues known to interact with SAM in the methyltransferase catalytic site of DOT1^102^ suggesting that it could be involved in both the methyltransferase and protease activities of DOT1. Notably, such dual transferase/protease function has been reported for the *O*-GlcNAc transferase (OGT).^103,104^ Interestingly, the C-terminus of DOT1L also has some similarity to a protein fold of certain serine proteases **(Supplemental Figure 6H & 6I)**.

To determine whether DOT1L has proteolytic activity, we obtained commercially available N-terminal DOT1L (abcam) and expressed the C-terminus of DOT1L in E. coli **(Figure 6G, yellow bar)**. We created in vitro cleavage reporter constructs with the protease-resistant HmuY protein as a scaffold attached to a region of c-MYC spanning MbII - MbIIIb, and MbIIIb - bHLH **(Figure 6H)**, which we observed to be cleaved in cells **(Figure 6E)**. The C-terminus of DOT1L was unable to cleave either region of c-MYC in vitro **(Figure 6I)**. On the contrary, the N-terminus of DOT1L cleaved the region between MbIIIb – bHLH in a dose-dependent manner **(Figure 6I, right panel)**. To determine whether in vitro cleavage of the MbIIIb – bHLH region of c-MYC can be recapitulated with cell extracts, we incubated the regions spanning MbII – IIIb, and IIIb – bHLH with either cytoplasmic or nuclear U2OS extracts for 24 hours at 37 °C. The nuclear extract cleaved the region spanning MbII – IIIb, while the cytoplasmic extract did not **(Figure 6J, left panel)**. Size-matching determined that in vitro cleavage of this c-MYC region is similar to the cleavage that occurs in the transfected HmuY construct **(Figure 6E, right panel)**. Both cytoplasmic and nuclear extracts led to efficient cleavage of the region between MbIIIb – bHLH with distinct banding patterns, indicating that cleavage of this c-MYC region can be facilitated by different proteases depending on c-MYC cellular localization **(Figure 6J, right panel)**. Cytoplasmic cleavage of c-MYC by calpains in this region was reported in the past,^99^ however the distinct nuclear extract-mediated cleavage has never been reported, to the best of our knowledge. Size matching of the nuclear cleavage at this c-MYC region determined that the site of in vitro cleavage is similar to that of the transfected nuclear HmuY construct containing this c-MYC region **(Figure 6E, right panel)**. Blotting for alpha-tubulin and histone 3 showed that cellular fractionation was successful in separating the two compartments **(Figure 6K)**.

Next, we incubated the full length commercially available recombinant c-MYC protein (abcam) with either the C-terminus or N-terminus of DOT1L to see whether DOT1L is capable of proteolytically cleaving it. Incubation with the N-terminus, but not the C-terminus, of DOT1L led to production of distinct ∼40 kDa N-terminal **(Figure 7A, upper panel)**, and corresponding ∼25 kDa C-terminal c-MYC bands **(Figure 7A, lower panel)**. When we incubated recombinant c-MYC with cytoplasmic, nuclear and chromatin extracts from U2OS cells, we observed a predominant N-terminal c-MYC cleavage band of ∼40 kDa with the nuclear extract **(Figure 7B, upper panel)** that matches in size to the cleavage product by N-terminal DOT1L. Similarly, probing for the C-terminal product revealed a ∼25kDa c-MYC fragment from the nuclear extract incubation **(Figure 7B, lower panel)** that matches the C-terminal product from DOT1L-mediated cleavage. Blotting for alpha-tubulin and histone 3 showed that cellular fractionation was successful in separating the three compartments **(Figure 6K & 7C)**.

**Figure 7.**
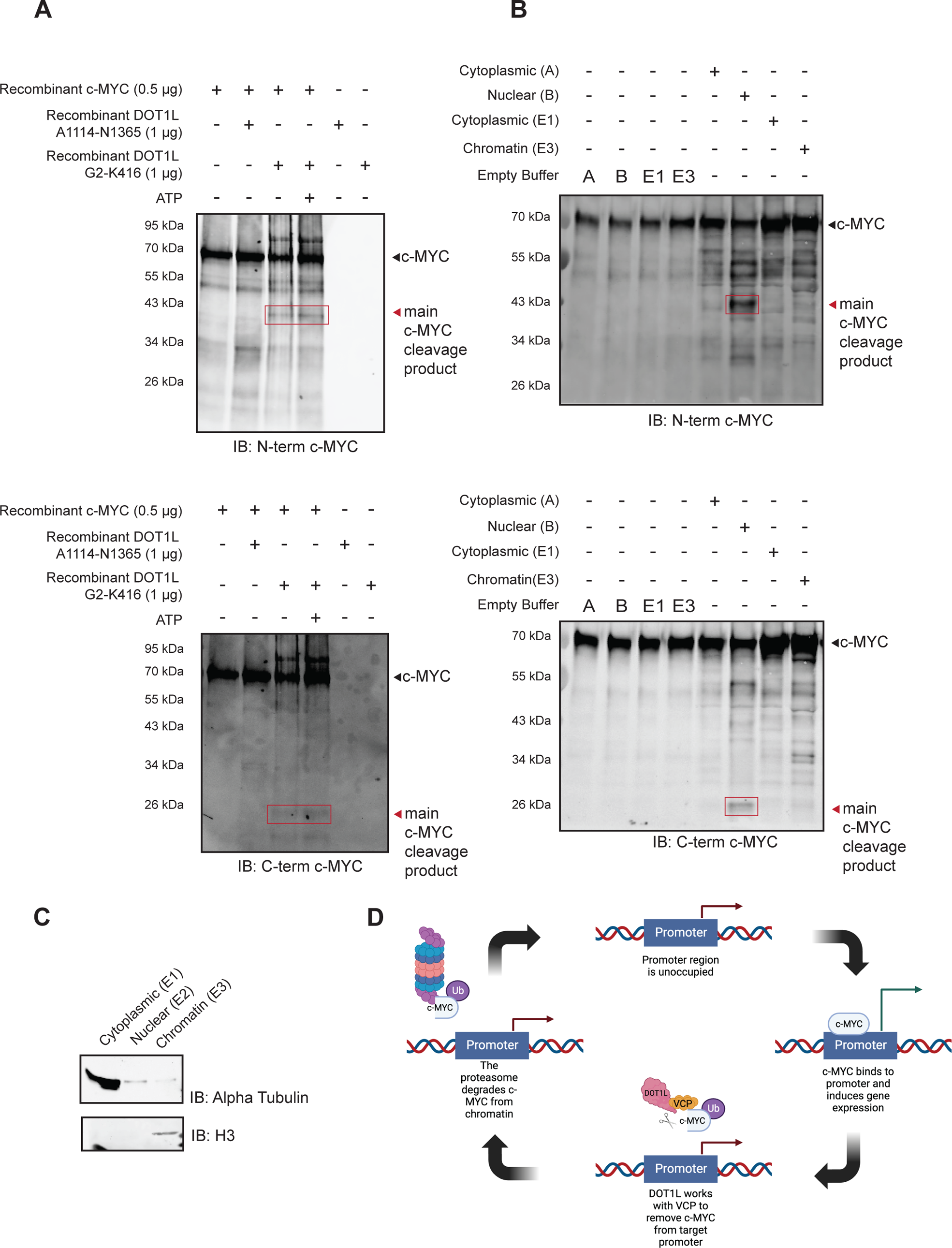
DOT1L cleaves full-length recombinant c-MYC in vitro. **A.** N- and C-terminal c-MYC cleavage fragments are produced after 24 hours of incubation with N-terminal DOT1L at 37 °C (N=2). Incubation with C-terminal DOT1L does not produce distinct cleavage fragments. **B.** Incubation of recombinant c-MYC with nuclear or chromatin fractions of U2OS cells produces N- and C-terminal cleavage fragments that match in size to the cleavage fragments produced by the N-terminus of DOT1L. **C.** Western blotting of alpha tubulin and H3 show that the cellular fractionation protocol successfully separated the cytoplasmic and chromatin fractions of U2OS cells. **D.** DOT1L works with the ubiquitin-proteasome system to displace c-MYC from target promoters. Unoccupied target promoters are bound by c-MYC. This induces the transcription of target genes and renders c-MYC as a “spent” transcription factor. DOT1L and VCP cooperate in the removal of c-MYC from the target promoter. DOT1L potentially acts by cleaving c-MYC, which may be needed for the removal of c-MYC by VCP. Displaced c-MYC is then degraded by the nuclear proteasome and allows new c-MYC to bind to the target promoter. Depletion of DOT1L disrupts the process of c-MYC displacement leading to prolonged occupancy of “spent” c-MYC, which then blocks new c-MYC from binding and inducing gene expression. This schematic was created with BioRender.com.

Overall, our findings are consistent with a model where DOT1L facilitates proper c-MYC turnover through the ubiquitin-proteasome system, particularly via its cooperation with VCP, and ensures accurate expression of c-MYC target genes. Our in vitro experiments show that DOT1L protease activity generates c-MYC fragments. We propose that the proteolytic cleavage of c-MYC is an important step in its turnover at chromatin in vivo, potentially by increasing recognition of the cleavage products by the proteasome system **(Figure 7D)**. Based on the presented results, we believe that the protease activity of DOT1L is conserved from *C. elegans* to mammals, although the precise mechanism of target protein cleavage by the DOT1L protease remains to be investigated.

## DISCUSSION

### DOT1L and transcription

DOT1L is the only known H3K79 methyltransferase in mammals^38,40,41^ and the major H3K79-methylating enzyme in *C. elegan**s***.^42^ While H3K79 methylation is associated with active gene transcription,^37^ the mechanism by which it affects transcription is not yet fully elucidated. Evidence from cultured cells indicates that DOT1L may either act with the super elongation complex (SEC) to influence transcriptional elongation^39^ or recruit TFIID, thus regulating transcription initiation.^45^ DOT1L was also reported to interact with RNA polymerase II to drive the expression of pluripotency markers.^105^ Moreover, we have previously reported that DOT-1.1 can negatively modulate the transcription of highly expressed target genes during postembryonic development in *C. elegans*.^42^ In this study, we describe a novel mechanism of transcription regulation by DOT1L relevant to the MYC transcription factor network. We show that DOT1L/DOT-1.1 and c-MYC/MML-1 co-regulate genes genome-wide in both *C. elegans* and in triple-negative breast cancer cells. We demonstrate that without DOT1L/DOT-1.1, inactive c-MYC/MML-1 accumulates at promoters and does not have competence in transcription regulation of its targets. Finally, we provide evidence that DOT1L acts in the same pathway with the ubiquitin-proteasome system, particularly with VCP, to regulate c-MYC occupancy on chromatin and, ultimately, c-MYC-driven transcription.

Our data are consistent with the HMT-independent role of DOT1L in c-MYC activation and specifically with the possibility that a novel protease activity of DOT1L might be involved. Notably, it was shown that catalytically inactive DOT1L can still regulate transcription and development,^41,42^ and methyltransferase-independent activity of the Dot1 protein was observed in yeast.^106^ Since the methyltransferase and protease domains of DOT1 enzymes overlap, it will be important to find amino acid residues whose mutation separates these activities. Moreover, our results do not exclude the possibility of the scaffold role of DOT1L that may be independent of both catalytic activities.

### c-MYC and DOT1L in cancer

DOT1L is a relatively new oncoprotein. However, the number of studies implicating DOT1L in human cancer is steadily growing. First, it was shown to promote mixed-lineage leukemia development through a mechanism whereby excessive H3K79 methylation at the HOXA9 and MEIS loci antagonizes repressive chromatin modifications normally present in differentiated blood cells.^107,108^ Chemical inhibitors of the DOT1L HMT activity are currently in clinical trials for treating leukemia.^34,109^ Recently, DOT1L has been recognized as a co-factor of the estrogen receptor in the breast,^110^ and its expression has been associated with ER-negative^27^ and triple-negative^28,111^ tumors. Additionally, high levels of DOT1L expression correlate with poor outcomes in ovarian^29,112–114^ and colorectal^23,115^ cancers, and alterations in the DOT1L gene have been linked to lung adenocarcinoma^116^ and pancreatic cancer.^117^ Since c-MYC amplification or overexpression is prevalent in cancer,^1^ our findings suggest that DOT1L could play a global role in controlling oncogenic c-MYC activity.

In several cases, the oncogenic function of DOT1L has already been linked to c-MYC activation.^38^ DOT1L was shown to cooperate with c-MYC in a metastatic breast cancer model,^28^ and a functional connection between DOT1L and MYCN has been reported in neuroblastoma.^35^ Most recently, a detailed clinical study correlated the severity of colorectal cancer disease with the level of DOT1L expression, which was highly elevated.^23^ Moreover, the authors demonstrated that c-MYC activation downstream of DOT1L was largely responsible for the oncogenic phenotypes resulting from DOT1L overexpression.^23^ Our data further implicates DOT1L in the global regulation of c-MYC-driven transcription.

### Transcription factor activation via the ubiquitin-proteasome system

The counterintuitive idea that a transcription factor needs to be degraded in order to activate gene transcription is not new. It was proposed that after a transcription factor binds and induces gene expression, it can exist in a “spent” state that needs to be removed for a new transcription factor molecule to bind to the promoter.^73^ The “spent” transcription factor would then need to be degraded, likely by the nuclear proteasome system.^72,118,119^ In yeast, GCN4, GAL4, and INO2/4 were shown to be dependent on proteasomal degradation for driving active gene transcription.^120^ Similar to our findings using *DOT1L KO/+* cells, despite an increase in GAL4 promoter occupancy upon MG132 treatment, a decrease in GAL4 target gene expression was observed.^120^ Another study showed that the recruitment of a transcription activator domain to DNA-bound receptor proteins increased their degradation such that the rate of degradation correlated with the potency of the activating domain, indicating that transcription activation and proteasomal degradation are linked.^121^

Furthermore, there are multiple transcription factors, including c-MYC, in which transactivating domains overlap with degrons signaling for proteasomal degradation.^122^ The DNA binding and transcriptional activity of these transcriptional activators are essential for their turnover.^121,123^ For example, a recent study showed that mutations in the conserved Myc Box II can lead to the accumulation of c-MYC at the *EZH2* promoter without an increase in *EZH2* gene expression, indicating that inactive c-MYC can accumulate at promoters and its transcriptional activity is needed for turnover.^124^

The importance of c-MYC proteasomal degradation for gene transcription was confirmed in a 2016 study showing that a K-less version of c-MYC cannot induce gene transcription since it is unable to be ubiquitinated and subsequently degraded.^18^ Furthermore, it was shown that inhibition of the proteasome led to increased c-MYC binding to promoters and that the degradation of c-MYC was needed for the transfer of PAF1C to RNA polymerase II, in order to drive active gene transcription.^18^ ^22^ In our study, we show a similar observation in *DOT1L KO/+* cell lines. When DOT1L is depleted, we see an accumulation of c-MYC at its target promoters, without a corresponding increase in target gene expression.

An important player in the ubiquitin-proteasome degradation system, VCP is a homohexameric AAA-ATPase that has important functions in ubiquitin-dependent protein homeostasis, most notably in endoplasmic reticulum-associated degradation (ERAD) and autophagy.^125^ Functionally, VCP acts primarily as a segragase and unfoldase to extract target proteins from membranes or protein complexes, while its role in the removal of chromatin-associated proteins has been recognized to be important in the DNA damage response pathway, DNA replication and mitosis.^126–128^ On the other hand, the role of VCP in transcription is slowly being recognized. During transcriptional stress, VCP is needed for the removal and degradation of RNA polymerase II at sites of transcriptional stalling.^129,130^ VCP also regulates the removal of the transcriptional repressor α2 from its target promoters^131^ and the degradation of transcription factor HIF1α.^132^ More recently, VCP has been implicated in the degradation and extraction of c-MYC from chromatin.^71^ Interestingly, VCP inhibition and depletion lead to a decrease in c-MYC target gene expression, highlighting the importance of c-MYC removal from target promoters.^71^

In our study, proteasomal and VCP inhibition led to a decrease in c-MYC target gene expression in the control triple-negative breast cancer cells, further validating the role of the ubiquitin-proteasome system in the activation of c-MYC target gene expression. This decrease is not seen in the *DOT1L KO/+* cell line under proteasomal and VCP inhibition conditions. Furthermore, DOT1L depletion exhibits slower dynamics of ubiquitinated protein clearance when the ubiquitin-activating enzyme is inhibited. Altogether, this highlights the importance of DOT1L in the turnover of c-MYC at chromatin.

### The role of proteolytic cleavage in c-MYC regulation

c-MYC regulation via a different type of proteolytic cleavage has been described.^99,133,134^ It occurs via calcium-activated calpain cleavage producing the N-terminal Myc-nick product.^99^ This cleavage product is transcriptionally inactive and is located in the cytoplasm, where it associates with α -tubulin and promotes α-tubulin acetylation by GCN5.^99,100^ Myc-nick has been implicated in cancer through its role in enhancing cancer cell survival and motility, which leads to tumor progression.^100,133^ In contrast to Myc-nick, we observe nuclear c-MYC fragments that arise from cleavage between Myc Box II and IIIb, as well as the region between Myc Box IIIb and the bHLH region, the latter of which is facilitated by DOT1L.

Surprisingly, the HMT domain of DOT1L has a similarity to the catalytic site of a class of aspartyl acid proteases known to recognize and cleave substrates with long ubiquitin chains.^92,93^ DDI-1 in particular has been shown to cleave ER-associated SKN-1A/NRF1 in *C. elegans* to activate the transcription factor in order for SKN-1A to mediate proteasome homeostasis.^84^ The mammalian homolog DDI-2 has also shown protease activity on NRF1 in mammalian cells.^91,92^ Notably, the activation of NRF1 is dependent on its extraction from the ER by VCP.^85^ Our data suggests that DOT1L-mediated cleavage of c-MYC may be important for its turnover at chromatin, and the continuous expression of its target genes.

Our data are consistent with a model wherein DOT1L proteolytically cleaves c-MYC bound to chromatin, thereby making it a target for the ubiquitin-proteasome system. VCP and DOT1L may work together in the removal of “spent” ubiquitin-modified c-MYC from chromatin, thus allowing newly produced c-MYC to bind promoters and activate transcription. Furthermore, cleavage of c-MYC may lead to N-end rule ubiquitination^87^ of the cleavage products, potentially making it a more attractive substrate of VCP. Of note, a close relative of NRF1, NFE2L2 (NRF2), exhibits downregulated pathway expression in the *DOT1L KO/+* cells compared to wild-type **(Figure 2D)**, indicating that the role of DOT1L on transcription factor turnover may extend beyond c-MYC.

### New avenues for targeting c-MYC in cancer

In addition to being a transcription factor, c-MYC has intrinsically disordered domains that make it a difficult target for small molecule inhibition.^135^ Currently, there are no direct c-MYC inhibitors. Instead, researchers have created drugs that target specific steps in *c-MYC* gene expression or MYC heterodimerization with MAX; additionally, a dominant negative mutant OmoMyc protein has been designed.^135–138^ Despite these efforts, the above approaches are not yet used in the clinic, and there exists a need to find new ways for inhibiting c-MYC activity. The mechanism of c-MYC activation that we uncovered, which involves a DOT1L-promoted c-MYC degradation on chromatin, represents a novel avenue for developing anti-cancer drugs. In the future, it might be possible to combine existing proteasome inhibitors used in the clinic with more specific inhibitors of DOT1L protease function for cancer treatment. This is especially exciting because overactive c-MYC is a feature of most human malignancies^6^ and targeting it would provide a broadly applicable approach.

### Limitations of the study

DOT1L is a large protein (1,739 aa). The methyltransferase activity of DOT1L is commonly studied in vitro using its N-terminal 400 aa domain and we assigned the novel protease activity to this same domain. However, the protease activity may be best revealed with the full-length DOT1L protein or its complex with interacting partners that may promote DOT1L protease activity by altering the domain conformation. We obtained evidence of DOT1L protease activity using commercially available DOT1L G2-K416 preparations (ab80254) which showed target cleavage patterns resembling that of nuclear extract preparations **(Figure 7)**. However, even the most active preps were rapidly losing their activity during storage, and preparations made in the laboratory had very modest cleavage capacity. Although we have preliminary data with the mutant DOT1L construct suggesting that mutation of the putative catalytic aspartic acid in the FW**D**LG sequence affects the efficiency of target cleavage, our inability to consistently obtain active wild-type DOT1L protease samples prevents us from reporting these data. Ultimately, DOT1L protease assays may require additional interacting partners as well as properly ubiquitin-modified cleavage substrates.

## Acknowledgments

The anti-MML-1 antiserum was a gift from Don Ayer (University of Utah). *Porphyromonas gingivalis* (strain 381) was obtained from Robert Berland and Caroline Genco (Tufts University). We thank Daniel Cifuentes, Nelson Lau, Mikel Garcia-Marcos, Matthew Layne, Mike Blower, Shawn Lyons, Shoumita Dasgupta, and their lab members for discussions and reagents. We thank Thomas Liontis for his help with *C. elegans* work and discussions, and Stefanie Chan for her help with cell culture for proteomics. We also thank the Boston University Genome Science Institute for RNA-seq funding. This research was financially supported by the National Institutes of Health (NIH) awards P30DK063608 and R01GM135199 to AG, the Schaefer Research Scholars Program Award to AG, the Dahod Breast Cancer Research Pilot Grant Award to AG and MDC, the Department of Defense Breast Cancer Breakthrough Award BC160363 to VP and MDC, the Ignition Award from the Office of Technology Development at Boston University (AG), and Boston University startup funds (AG). Strains were provided by the *C. elegans* Gene Knockout Project at the Oklahoma Medical Research Foundation and by the *C. elegans* Reverse Genetics Core Facility at the University of British Columbia, which are part of the International *C. elegans* Gene Knockout Consortium. Some strains used in this study were obtained from the *Caenorhabditis Genetics Center*, which is funded by the NIH Office of Research Infrastructure Programs (P40OD010440). The BU Microarray and Sequencing Resource Core Facility is supported by the UL1TR001430 grant awarded to the BU Clinical and Translational Science Institute.

## Author Contributions

Conceptualization, A.G.; Methodology, G.P.S., E.S.G., A.M., R.E., A.C., J.K., and M.D.C.; Investigation, G.P.S., R.E., E.S.G., M.D.C., A.M., I.N., A.C., P.D., J.K., and B.C.B.; Writing – Original Draft, G.P.S., and A.G.; Writing – Review & Editing, G.P.S., and A.G..; Funding Acquisition, A.G., M.D.C., V.P., and A.E.; Resources, A.G., V.P., and A.E.; Supervision, A.G., V.P., M.D.C., and A.E.

## Conflict of interest

The authors declare no competing interests.

---

**Supplemental Figure 1.**
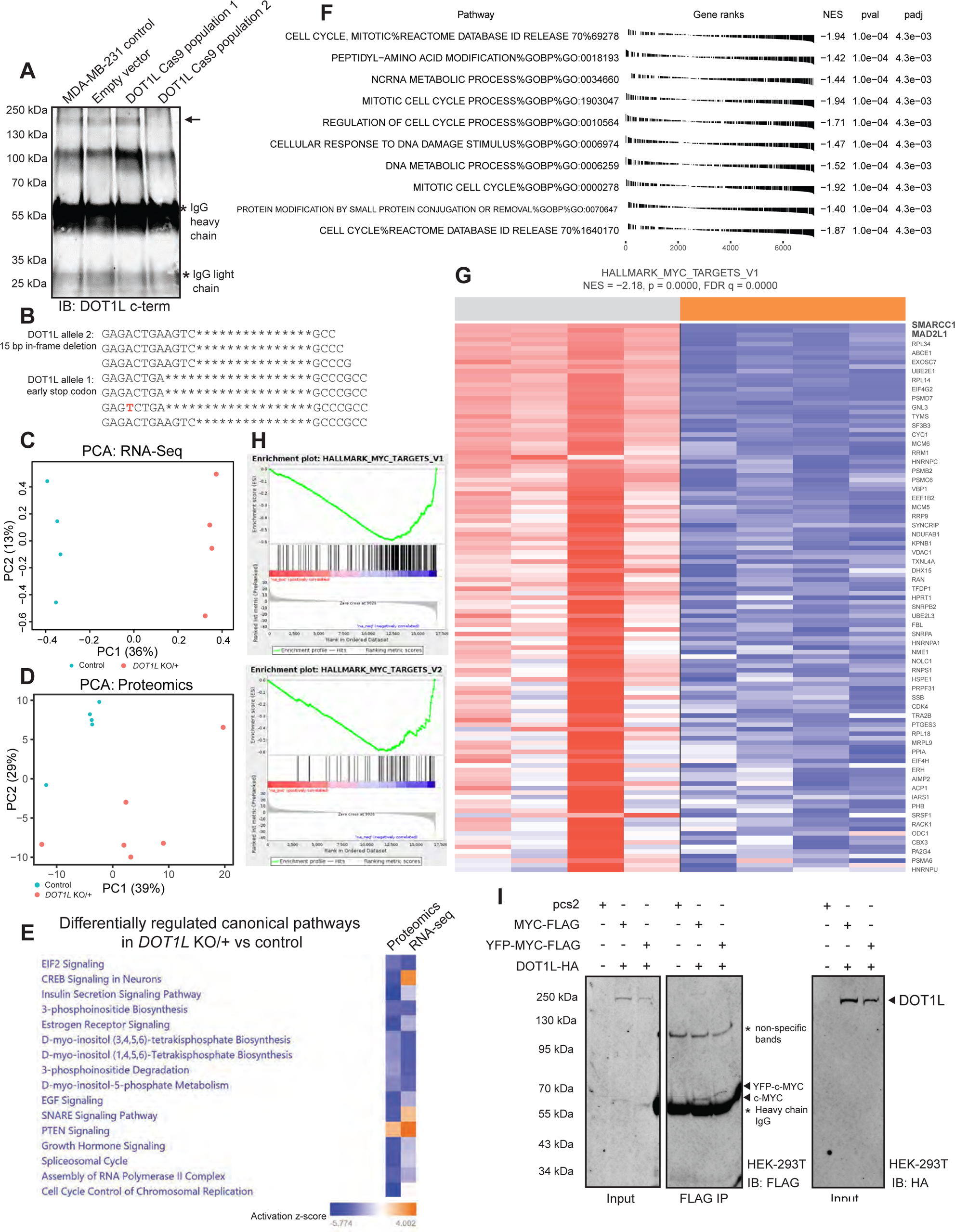
Bulk RNA-seq and proteomic analyses show that the c-MYC pathway and hallmark c-MYC targets are downregulated in the *DOT1L KO/+* cells compared to the control cells. **A.** IP-western blot of two MDA-MB-231 populations after CRISPR/Cas9 gene editing to knock out DOT1L. The second population (fourth lane) showed successful knockout of the full-length DOT1L protein and was then used to isolate a clonal *DOT1L KO/+* line. **B.** RNA-seq reads of the DOT1L locus show that there are two alleles present in the *DOT1L KO/+* cells. There is a 19-bp deletion leading to an early stop codon, and a 15-bp in-frame deletion. **C.** Principal Component Analysis of RNA-seq data shows that the wild-type and *DOT1L KO/+* cells primarily group with their respective genotype (N=4). **D.** Principal Component Analysis of proteomic data shows that the wild-type population and *DOT1L KO* primarily group with their respective genotype (N=5). **E.** Top differentially regulated canonical pathways in the *DOT1L KO/+* MDA-MB-231 cells compared to the wild-type counterpart as analyzed by Ingenuity Pathway Analysis. **F.** Gene set enrichment analysis (GSEA) of proteomic data shows that the cell cycle and protein modification pathways are among the most downregulated in the *DOT1L* KO population. **G.** Heatmap shows that c-MYC hallmark targets gene expression is downregulated in *DOT1L KO/+* cells. **H.** Gene set enrichment analysis (GSEA) plots from RNA-seq analysis show that c-MYC Hallmark genes are downregulated in *DOT1L KO/+* cells. **I.** Western blot showing successful immunoprecipitation of overexpressed MYC and YFP-MYC in HEK-293T cells.

**Supplemental Figure 2.**
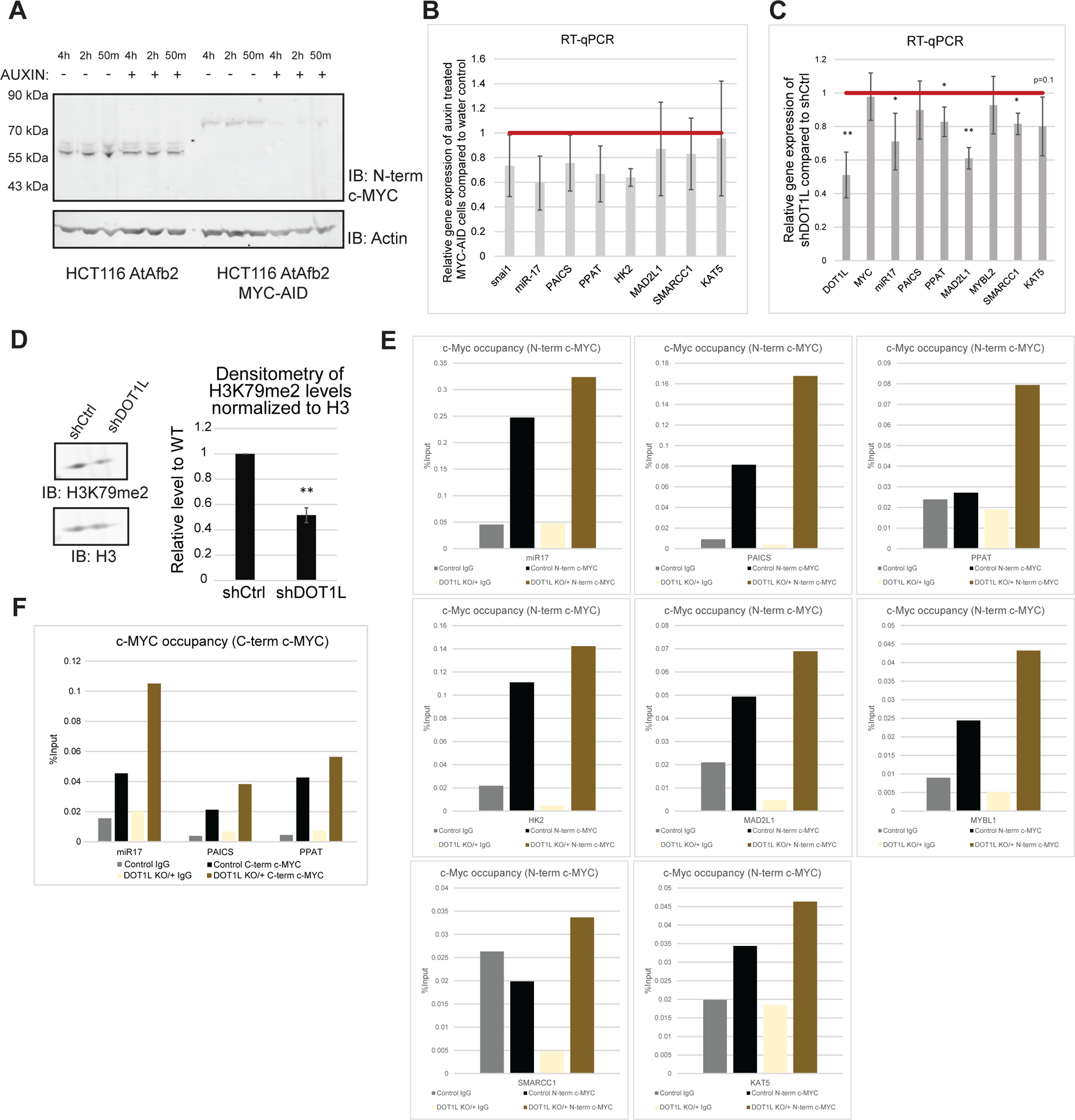
DOT1L depletion leads to a decrease in c-MYC target gene expression, but an increase in c-MYC occupancy at the corresponding promoters. **A.** Endogenous c-MYC tagged with the auxin-inducible degron (AID) is depleted after 50 minutes of auxin treatment (0.1 mg/mL) (N=3). Endogenous c-MYC without the AID in the HCT116 cells containing the auxin receptor F-box protein AtAFB2 is not depleted when treated with auxin. **B.** Auxin treatment for 4 hours leads to a decrease in c-MYC target gene expression via RT-qPCR (N=3). **C.** shRNA against DOT1L in HCT116 cells leads to a decrease in c-MYC target gene expression (N=4). **D.** shRNA against DOT1L in HCT116 cells leads to a decrease in H3K79me2 (N=4) **E.** A representative biological replicate of c-MYC ChIP-qPCR using the N-terminal c-MYC antibody (Y69) showing that c-MYC occupancy is increased at its target gene promoters in *DOT1L KO/+* cells compared to wild-type (N=4). **F.** A representative biological replicate of c-MYC ChIP-qPCR using the C-terminal c-MYC antibody (9E10) showing that c-MYC occupancy is increased at its target gene promoters in *DOT1L KO/+* cells compared to wild-type (N=3). Asterisks *, ** and *** denote P values (Student’s t-test) less than 0.05, 0.01 and 0.001, respectively.

**Supplemental Figure 3.**
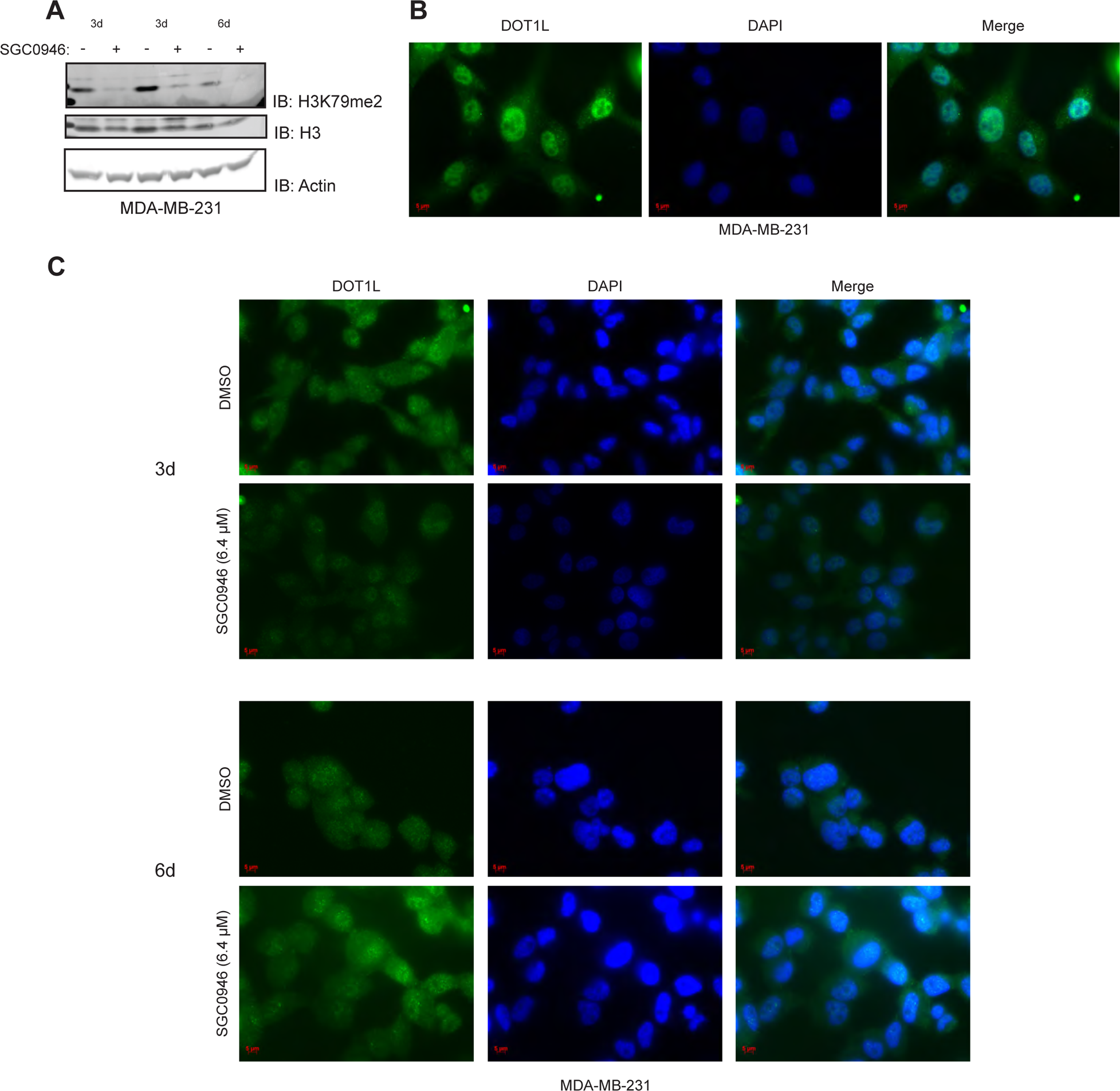
Inhibition of the HMT activity of DOT1L does not alter DOT1L localization. **A.** Inhibition of HMT activity by SGC0946 (6.4 μM) for 3 (N=2) or 6 days (N=1) leads to the depletion of H3K79me2 in MDA-MB-231 cells. **B.** Immunocytochemistry for DOT1L (ab64077) in untreated MDA-MB-231 cells shows the nuclear localization of DOT1L. **C.** Inhibition of HMT activity by SGC0946 (6.4 μM) for 3 (N=2) or 6 days (N=1) does not lead to changes in DOT1L nuclear localization.

**Supplemental Figure 4.**
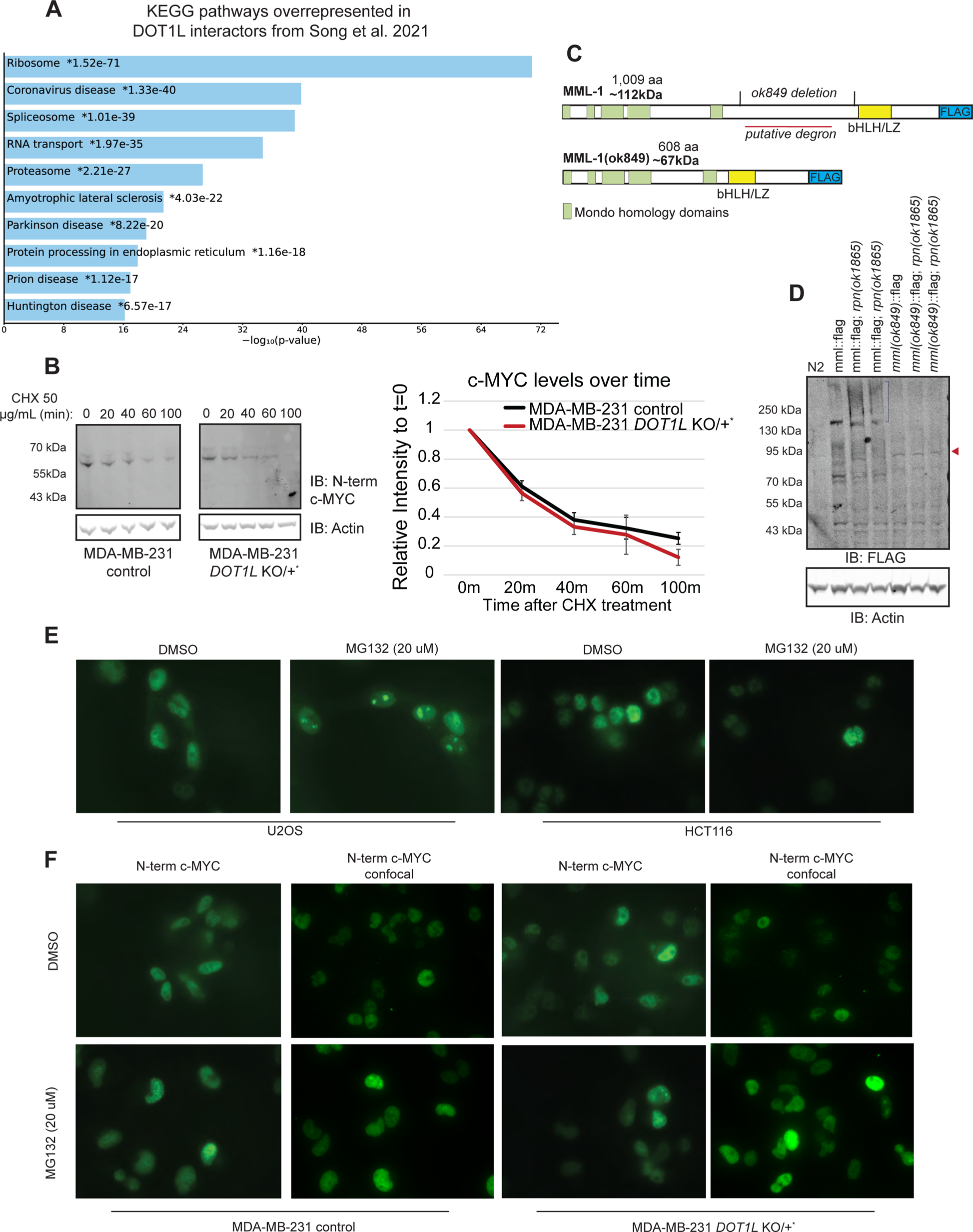
DOT1L works with the ubiquitin-proteasome system to regulate c-MYC. **A.** KEGG pathway analysis by EnrichR shows that DOT1L binding partners^33^ are enriched in the ribosome, spliceosome, and proteasome components. **B.** Cycloheximide (CHX) chase assays show that overall c-MYC stability is minimally decreased in *DOT1L KO/+* cells compared to control (N=3). **C.** A schematic of the FLAG-tagged *C. elegans* MML-1 protein and the MML-1(ok849) protein produced in the *mml-1(ok849)* mutant. **D.** Endogenously FLAG-tagged MML-1 is stabilized in the *rpn-10(ok1865)* worms as evidenced by the smear above the full-length MML-1 protein (N=2). This smearing is not observed with the inactive MML-1(ok849) mutant in the same *rpn-10* mutant background. **E.** 4-hour treatment with 20 μM MG132 leads to the formation of c-MYC multimers in U2OS and HCT116 cells (N=2). **F.** *DOT1L KO/+* cells do not readily form c-MYC multimers compared to the wild-type controls. Both the control and *DOT1L KO/+* cells form c-MYC multimers upon 4-hour treatment with 20 μM MG132 (N=2).

**Supplemental Figure 5.**
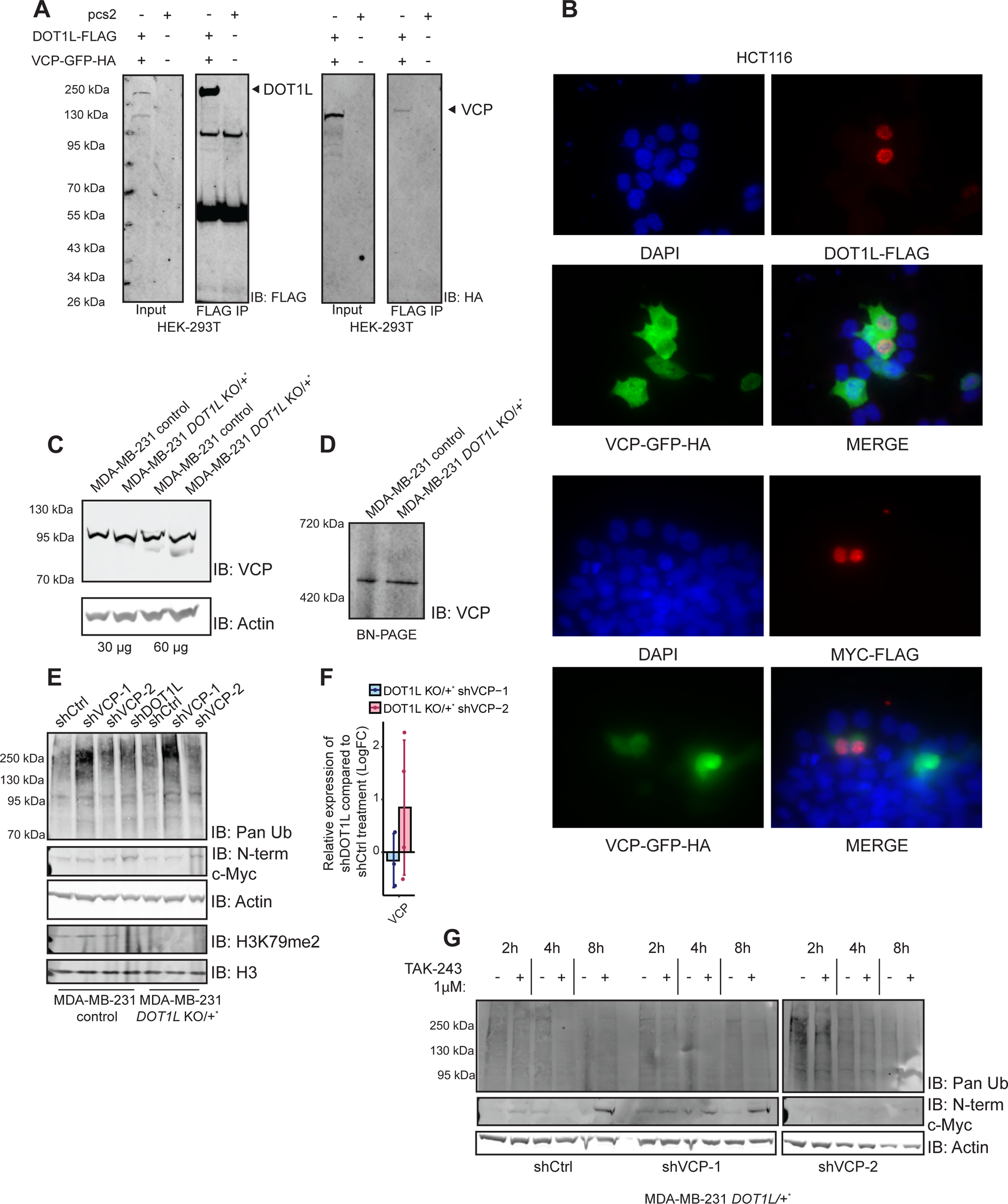
VCP and DOT1L cooperate to regulate c-MYC-driven gene transcription. **A.** Overexpressed VCP-GFP immunoprecipitates with overexpressed DOT1L in HEK-293T cells (N=2). **B.** Overexpressed VCP-GFP localizes to the cytoplasm and the nucleus, while over expressed DOT1L and c-MYC localize to the nucleus only in HCT116 cells. **C.** DOT1L depletion does not alter VCP protein levels. **D.** DOT1L depletion does not alter hexamerization of VCP. Control and *DOT1L KO/+* lysates were run on a BN-PAGE gel to see native protein complexes. **E.** Both shRNAs against VCP leads to an increase in ubiquitinated proteins in the control MDA-MB-231 cells, while only shVCP-1 led to an increase in ubiquitinated proteins in *DOT1L KO/+* cells. shDOT1L leads to a decrease in H3K79me2 in control MDA-MB-231 cells. **F.** shVCP-1 leads to a slight decrease in VCP mRNA expression in *DOT1L KO/+* cells, while shVCP-2 does not. **G.** TAK-243 chase experiments show that proteasomal clearing of ubiquitinated substrates is unchanged in its delay between shCtrl and shVCP *DOT1L KO/+* MDA-MB-231 cells.

**Supplemental Figure 6.**
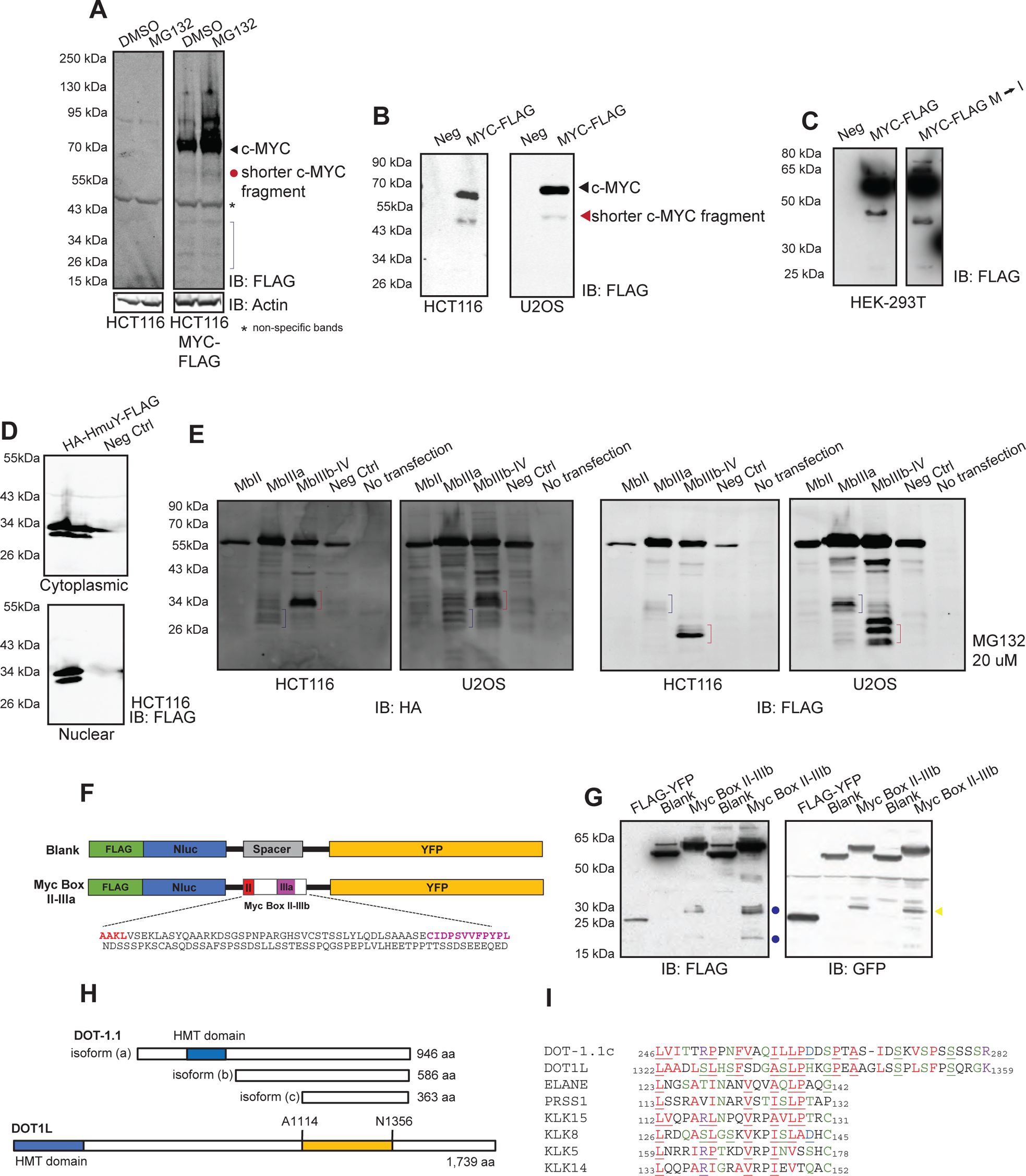
Shorter c-MYC fragments are observed endogenously and through reporters. **A.** Endogenously FLAG-tagged c-MYC in HCT116 cells show evidence of shorter c-MYC fragments that are stabilized when treated with the proteasome inhibitor MG132 (N=3). **B.** Transfection of FLAG-tagged c-MYC in HCT116 (N=3) and U2OS (N=4) cells consistently produces a ∼45 kDa c-MYC fragment. **C.** The 45 kDa c-MYC fragment produced from c-MYC transfections is not a result of an internal start codon (N=3). Transfection of a c-MYC-FLAG construct with the internal Methionine residues mutated to Isoleucine still produced the 45 kDa c-MYC fragment. **D.** The HmuY protein from *Porphyromonas gingivalis* can be expressed in HCT116 (N=1) cells. The HmuY protein is expressed in both the cytoplasm and nucleus. **E.** Transfection of the HmuY constructs shows cleavage of c-MYC at the 143-210 aa region, and 258-320 aa region, as detected by both the HA and FLAG antibodies. These cleavage products are stabilized by proteasomal inhibition with 20 μM MG132. **F.** A schematic of the nanoluciferase-YFP reporters used to detect proteolytic cleavage in the middle portion of c-MYC. Either a random spacer sequence or amino acids 141-258 of c-MYC was inserted in between the nanoluciferase and YFP construct. **G.** Western blot using the FLAG and GFP antibodies shows N-terminal (blue dots) and C-terminal (yellow arrow) cleavage fragments arising from the middle portion of c-MYC. The N-terminal fragments have sizes of around 20 kDa and 27.5 kDa, while the C-terminal fragment detected has a size of around 35 kDa (N=3). **H.** A schematic of DOT-1.1 isoforms (*C. elegans*) and DOT1L (mammalian) protein. Highlighted in yellow is a region in the C-terminus that has homology to serine proteases. **I.** CLUSTAL Omega alignment of DOT-1.1c and the C-terminus of DOT1L shows conservation of a protein fold of serine proteases.

**Supplemental Table 1. Total RNA sequencing results comparing control (4) and clonal *DOT1L* KO/+ (4) MDA-MB-231 samples**

**Supplemental Table 2. Proteomics and phosphoproteomics results comparing control (5) and population #2 *DOT1L* KO (6) MDA-MB-231 samples**

## Experimental Procedures

### *C. elegans* strains and growth

All *C. elegans* strains were maintained at 20 °C using standard methods.^139^ The wild-type strain used was the *C. elegans* Bristol strain N2. The following mutant strains were obtained from the Caenorhabditis Genetics Center (CGC): VC20674, containing *dot-1.1(gk105059)*I, VC20787, containing *mml-1(gk402844)*III, RB954 - *mml-1*(ok849)III, VC1369 - *rpn-10(ok1865)*, RB653 - *ogt-1(ok430)*III, RB774 - *zfp-1(ok554)*III, and OP198 - *unc-119(ed3)*III; wgIs198[MML-1::TY1::EGFP::3xFLAG(92C12) + unc-119(+)]X. The RB and VC strains were provided by the *C. elegans* Gene Knockout Project at the Oklahoma Medical Research Foundation and by the *C. elegans* Reverse Genetics Core Facility at the University of British Columbia, respectively, which were part of the International *C. elegans* Gene Knockout Consortium. The MML::FLAG and MML(ok849)::FLAG endogenous locus tagging was done by InVivo Biosystems via CRISPR-Cas9. We isolated homozygous tagged lines. Mutations of interest were outcrossed four times and the following derivative strains were obtained: AGK760 - *mml-1(gk402844)*III, AGK784 - *dot-1.1(gk105059)*I, AGK773 - *dot-1.1(gk105059)*I; *mml-1(gk402844)*III, AGK776 - *ogt-1(ok430)*III; wgIs198X, AGK744 - *zfp-1(ok554) unc-119(ed3)*III; wgls198X, AGK878 - *rpn-10(ok1865)*; *mml-1*::FLAG, AGK911 - *rpn-10(ok1865)*; *mml-1 (ok849)*::FLAG.

### RNA extraction and quantitative RT-PCR (*C. elegans*)

Hypochlorite-synchronized L1 stage wild type or mutant animals were grown for 36 hours post-hatching at 20°C on standard NGM plates with OP-50 E. coli. Animals were washed off the plates using isotonic M9 solution. RNA was extracted using TRIzol (Invitrogen) followed by miRNeasy mini column (Qiagen). cDNA was generated from 1-2 mg of total RNA using random hexamers and Maxima Reverse Transcriptase (Thermo Fisher Scientific). Quantitative PCR was performed on the ViiA 7 Real-Time PCR System (Life Technologies) using the QuantiFast SYBR Green PCR Kit (Qiagen). Thermocycling was done for 40 cycles in a two-step cycling in accordance with the manufacturer’s instructions and each PCR reaction was performed in triplicate. Changes in mRNA expression were quantified using *act-3* mRNA as a reference.

### *C. elegans* expression profiling

RNA was isolated in triplicate for each mutant strain, and its integrity was confirmed using a 2100 Bioanalyzer (Agilent Technologies). Samples were labeled and hybridized to Affymetrix GeneChip™ C. elegans Gene 1.0 ST arrays. Raw Affymetrix CEL files were normalized to produce gene-level expression values using the Robust Multiarray Average (RMA)^140^ in the Affymetrix Expression Console (version 1.4.1.46). The default probe sets defined by Affymetrix were used to assess array quality using the Area Under the [Receiver Operating Characteristics] Curve (AUC) metric. Principal Component Analysis (PCA) was performed using the prcomp R function with expression values that had been normalized across all samples to a mean of zero and a standard deviation of one. Differential expression was assessed using the moderated (empirical Bayesian) ANOVA and t test implemented in the limma package (version 3.14.4) (i.e., creating simple linear models with lmFit, followed by empirical Bayesian adjustment with eBayes). Correction for multiple hypothesis testing was accomplished using the Benjamini-Hochberg false discovery rate (FDR).^141^ All microarray analyses were performed using the R environment for statistical computing (version 2.15.1).

### Cell culture, transfection and protein extraction

The MDA-MB-231, and HCT116 human cells (ATCC Cell Bank) were grown in high-glucose DMEM medium supplemented with 10% FBS. For DOT1L inhibitor experiments, cells were treated with 6.4 uM SGC0946 (SelleckChem) for 3 days before lysis or fixation. For cycloheximide chase experiments, cells were treated with 50 μg/mL cycloheximide (Sigma, C7698) over a span of 2 hours before lysis and western blot. For proteasome experiments, cells were treated with 20 μM MG132 (Sigma) for 4 hours or 10 μM NMS-873 (Cayman Chemical) for 6 hours before lysis or fixation.

For transfection with plasmids, Lipofectamine 3000 was used according to the manufacturer’s protocol (Invitrogen). The following plasmids were transfected into U2OS, HEK-293T and HCT cells: pcs2, pcs2 HA-MYC-FLAG, pcs2 MYC-FLAG, pcs2 DOT1L-FLAG, pcs2 DOT1L-HA, pcs2 YFP-MYC-FLAG, the HmuY cleavage constructs, and VCP(wt)-EGFP, a gift from Nico Dantuma (Addgene plasmid # 23971; http://n2t.net/addgene:23971; RRID:Addgene_23971).^142^

The transfection of a mixture of specific siRNA against c-Myc protein (exon 2 and exon 3, IDs s9129 and s9130, respectively; Ambion) and DOT1L protein (IDs s39011, s39010; Ambion) was performed using Lipofectamine 3000 according to manufacturer’s protocol. The cells at 90% confluency were washed with 1xDPBS, treated with Trypsin 0.25% at 37 °C for 3 minutes, detached in 10 mL of complete DMEM and split onto 10cm plates 1:5. The transfection mixture was added to plates with the final concentration of siRNA equal to 10nM. The control cells were treated with the same concentration of the negative control silencer (Ambion, Cat. #AM4611). The protein extracts and the RNA samples were collected after 48 hours.

For the protein extraction, the cells were rinsed with cold 1xDPBS, scraped in 1xDPBS and centrifuged at 1,000 x *g* for 5 minutes at 4 °C. The pellet was then re-suspended in 200 μL of IPH lysis buffer (50 mM Tris-HCl pH 8.0, 250 mM NaCl, 5 mM EDTA, 1% NP-40 supplemented with fresh 0.1 mM PMSF and 1x Protease Inhibitor cocktail, EDTA-free (Thermo Scientific)) or RIPA lysis buffer (0.1% SDS, 1% Triton X-100, 0.1% Deoxycholate, 1 mM EDTA, 150 mM NaCl, 10 mM Tris pH 8.0 supplemented with 1x Protease Inhibitor cocktail and 1X NEM when applicable) and incubated for 20 minutes on ice. After that, the sample was centrifuged for 10 minutes at 21,000 x *g* at 4°C and the supernatant was collected.

For cell proliferation experiments, 0.1×10^6^ cells were plated on a twelve-well plate. Cells were collected for cell counting via the Countess II automatic cell counter at 24, 48 and 72 hours after plating. Cell counting experiments were done in triplicate.

### Generation of DOT1L KO cells

DOT1L knockout cells were generated using CRISPR-mediated genome editing of dot1l gene locus. The synthetic sgRNA oligo pair complementary to exon 1 was designed and cloned into lentiCRISPRv2 vector (Addgene #52961, gift from Dr. Feng Zhang)^143^ following the published protocol.^49^ For virus production, HEK293T cells were transfected with 8 μg lentiCRISPR-v2-DOT1LgRNA exon1 or lenti-CRISPR-v2 (empty vector control), 8 μg psPAX2 and 8μg VSV-G plasmids using Lipofectamine 3000. After incubation for 48 or 72 hours, the supernatants of the transfected cells containing viral vectors were collected and stored at – 80 °C. MDA-MB-231 cells were infected with the lentivirus encoding CRISPR/Cas9-DOT1LgRNA exon 1 or lentiCRISPR-v2 vector (control) at different concentrations. After 48 hours, the medium containing the lentivirus was replaced with high-glucose DMEM supplemented with 10% FBS and 2.5μg/mL of puromycin for selection of cells transduced with a lentiCRISPR construct. After a week, the populations of positive cells were collected and screened by western blot for the absence/presence of DOT1L protein.

### Generation of MYC-AID and MYC-FLAG cells

The E3 ligase AtAFB2 was inserted into the AAVS1 locus of HCT116 cells via CRISPR-Cas9 genome editing using the AAVS1 T2 CRIPR in pX330 vector, a gift from Masato Kanemaki (Addgene plasmid # 72833; http://n2t.net/addgene:72833; RRID:Addgene_72833)^65^ and the pSH-EFIRES-P-AtAFB2-mCherry-weak NLS vector as a donor, a gift from Elina Ikonen (Addgene plasmid # 129717; http://n2t.net/addgene:129717; RRID:Addgene_129717).^64^

HCT116 MYC-AID and MYC-FLAG cell lines were generated using CRISPR-Cas9-mediated genome editing of the *MYC* gene locus. sgRNA was generated complementary to the last exon of MYC and cloned into the pX330-U6-Chimeric_BB-CBh-hSpCas9 vector, a gift from Feng Zhang (Addgene plasmid # 42230; http://n2t.net/addgene:42230; RRID:Addgene_42230).^144^ For the MYC-AID lines, a donor pcs2 plasmid containing 200 base pairs flanking the stop codon of the *MYC* locus, a V5 tag, the mini IAA7 degron, a P2A self-cleaving peptide and the blasticidin resistance gene was co-transfected. Homozygous clones were isolated after two weeks of blasticidin treatment via serial dilution. The genotypes were confirmed using genomic PCR primers flanking the cut site and sanger sequencing, as well as western blotting. Auxin-induced degradation of c-MYC was tested using treatment with auxin at certain timepoints and checking via western blot for c-MYC protein levels.

For the MYC-FLAG cell lines, a donor pcs2 plasmid containing 200 base pairs flanking the stop codon of the *MYC* locus, 2x FLAG tags, a V5 tag, a P2A self-cleaving peptide and the blasticidin resistance gene was co-transfected. Homozygous clones were isolated after two weeks of blasticidin treatment via serial dilution. The genotypes were confirmed using genomic PCR primers flanking the cut site and sanger sequencing, as well as western blotting.

### shRNA knockdown

An shRNA scramble (CCTAAGGTTAAGTCGCCCTCG), a gift from David Sabatini (Addgene plasmid # 1864; http://n2t.net/addgene:1864; RRID:Addgene_1864),^145^ shRNA against DOT1L (CGCCAACACGAGTGTTATATT) and two shRNA sequences against VCP (CCTATCAACAGCCATTCTCAA, GATGGATGAATTGCAGTTGTT) were cloned into the pLKO.1 – Hygro vector, a gift from Bob Weinberg (Addgene plasmid # 24150; http://n2t.net/addgene:24150; RRID:Addgene_24150). The shRNA was co-transfected into HEK-293FT cells with 4 μg psPAX2 and 4 μg VSV-G plasmids using Lipofectamine 3000. The viral supernatant was collected 48 hours after transfection. The virus was then concentrated using PEG-8000 and resuspended in DPBS. In brief, 1 volume of sterile concentrator solution (40% PEG-8000, 1.2 M NaCl in PBS pH 7.4) was added to 3 volumes of cleared viral supernatant and incubated for at least 4 hours at 4 °C. The solution was then centrifuged at 1600xg for 60 minutes and resuspended in DPBS at 1/10 of the original volume.

Concentrated lentivirus containing the shRNA was added to HCT116 cells plated to 80-90% confluency. After 24 hours of infection, the cells were treated with hygromycin for two weeks before the population was collected for western blotting and RNA extraction.

### Chromatin Fractionation

Chromatin fractionation of MDA-MB-231 and U2OS cells was adapted from the fractionation protocol developed in 2018.^146^ Briefly, cells were washed with 1x PBS, scraped gently into a 1.5 mL microcentrifuge tube and pelleted at 130 x g at 4 °C. The cytoplasmic fraction was obtained by lysing the cells in 5x volume of Buffer E1 (50 mM HEPES-KOH, 140 mM NaCl, 1 mM EDTA, 10% glycerol, 0.5% NP-40, 0.25% Triton X-100, 1 mM DTT, 1x Protease Inhibitor Cocktail) pipetting up and down five times before pelleting at 1100 x g at 4 °C. The pellet was cleared of cytoplasmic lysate by repeating lysis in Buffer E1 twice more. The nuclear fraction was obtained by lysing the cells in 2x volume of Buffer E2 (10 mM Tris-HCl pH 8.0, 200 mM NaCl, 1 mM EDTA, 0.5 mM EGTA, 1x Protease Inhibitor Cocktail) pipetting up and down five times before pelleting at 1100 x g at 4 °C. The pellet was cleared of nuclear lysate by repeating lysis in Buffer E2 twice more. The chromatin fraction was obtained by lysing the cells in 5x volume of Buffer E3 (500 mM Tris-HCl pH 8.0, 500 mM NaCl) pipetting up and down five times. The chromatin fraction was then sonicated (Diagenode) on the high setting for 5 minutes (30s on, 1 min off).

### Nuclear Fractionation

Nuclear fractionation of U2OS cells was adapted from a previously described protocol.^99^ In brief, the cells were lysed with 5x pellet volume of Buffer A (10 mM HEPES pH 7.9, 10 mM KCl, 1.5 mM MgCl2, 0.5% NP40) for 20 minutes on ice. The cellular fraction was centrifuged at max rpm (15,000) and the supernatant was collected. The nuclear pellet was then washed twice more with Buffer A. The nuclear fraction was lysed in 4x volume of Buffer B (20 mM HEPES pH 7.9, 400 mM NaCl, 1.5 mM MgCl2, 0.2 mM EDTA, 15% glycerol) for 20 minutes on ice.

### Immunoprecipitation and western blotting

For immunoprecipitation, either 200-300 μg of cleared protein extract lysed in RIPA buffer was incubated with anti-FLAG M2 Magnetic Beads (Sigma) for 1 hour at 4 °C with rotation followed by 3 washes with RIPA buffer or 200-300 μg or protein extract was incubated with 2-5 μg of Y69 N-terminal c-MYC, FLAG M2 or HA-tag antibody at 4 °C overnight before precipitated with magnetic Protein G beads (Thermofisher) for 2 hours at 4 °C followed by 3 washes with RIPA buffer.

For co-immunoprecipitation, cells were lysed in co-immunoprecipitation buffer (20 mM HEPES pH 7.9, 200 mM NaCl, 0.5 mM EDTA, 10% glycerol, 0.2% NP-40) with 1x protease inhibitor cocktail and 1x phosphatase inhibitor (PhosSTOP, Roche) for 30 minutes on ice. The lysates were cleared via centrifugation at max rpm at 4 °C for 10 minutes. 50 μL of Protein G beads was preincubated with 2 μg of antibody at RT for 15-45 minutes before incubating with 500-1000 μg of protein lysates at 4 °C for 3-4 hours, followed 3 washed with co-immunoprecipitation buffer.

The precipitated proteins were extracted with 20 μL of 50 mM Glycine pH 2.8 and 10 μL of loading buffer (NuPAGE™ LDS Sample Buffer (4X) and NuPAGE™ Sample Reducing Agent (10X), Novex) at 70 °C for 10 minutes and used for the Western Blot. The input protein concentration was quantified by the Bradford assay. For C. elegans western blots, a single 60 mm^2^ plate of worms was collected and lysed directly in worm lysis buffer with 10% BME.

Proteins were resolved on precast NuPAGE Novex 4–12% Bis-Tris gels (Invitrogen), Bolt 8% Bis-Tris Plus gels (Invitrogen) or self-cast 10% acrylamide gels (0.375 M Tris pH 8.8, 0.1% SDS, 0.05% APS, 0.3% TEMED, 10% Acrylamide:Bis Acrylamide) for 1 hour and transferred to a nitrocellulose membrane (0.45µm) by semidry transfer (BioRad Trans-Blot SD transfer cell) at a constant current for 1 hour, or to a PVDF membrane (0.45 μm) by wet transfer overnight at a constant 7V overnight. The blotted membranes were blocked with the blocking buffer (5% non-fat dry milk in PBS-T) at room temperature for 30 minutes. Subsequently, they were incubated with appropriate primary antibodies overnight at 4 °C and with secondary antibodies for 1 hour at room temperature. Three washes with PBS-T buffer were made between and after the incubation with the antibodies. The membranes were visualized with the Odyssey Imaging System (LI-COR Biosciences, IRDye 800CW Goat anti-Rabbit IgG Secondary Antibody & IRDye 800CW Goat anti-Mouse IgG Secondary Antibody) or developed with SuperSignal West Femto kit (ThermoFisher) and scanned with KwikQuant Imager (KindleBiosciences). Antibodies against C-terminal (Abcam, 9E10, 1:2000) and N-terminal (Abcam, Y69, 1:1000) domains of MYC protein were used. Tagged proteins were detected by FLAG-HRP conjugated antibody (Sigma, A8592, 1:4000), mouse HA-probe antibody (Thermofisher, 2-2.2.14, 1:2000), rabbit HA-probe antibody (CST, C29F4, 1:1000), FLAG M2 Mouse Antibody (Sigma-Aldrich, 1:2000), and GFP rabbit Antibody (Abcam, ab290, 1:5000). Endogenous proteins were detected by rabbit DOT1L rabbit antibody (Abcam, ab64077, 1:500), VCP antibody (Abcam, EPR3308, 1:5000), mouse alpha tubulin antibody (house-made, E-74, 1:1000), rabbit GAPDH antibody (Abclonal, A10868, 1:1000), rabbit H3 antibody (Millipore, AS3, 1:2000), rabbit H3K79me2 antibody (Millipore, NL59, 1:1000), mouse Pan-ubiquitin antibody (CST, PD41, 1:1000), and mouse Actin antibody (Sigma, C4, 1:5000).

### Immunocytochemistry

Cells were directly plated onto coverslips treated with Poly-D-Lysine (Sigma) and grown overnight to around 80% confluency. The cells were fixed for 10 minutes with 4% formaldehyde (Fisher), permeabilized for 30 minutes with 0.25% Triton-X-100 and blocked for 30 minutes with 3% BSA in PBS. The fixed cells were incubated with the following primary antibodies at 4 °C overnight: mouse FLAG M2 antibody (1:200), rabbit HA antibody (1:200), rabbit VCP (1:200), DOT1L (1:200), N-terminal c-MYC (1:200). The fixed cells were then washed three times with PBS before incubating with the corresponding secondary antibodies at room temperature for 1 hour. After washing three times, the fixed cells were stained with DAPI (1:10,000) for 5 minutes and then washed three times with PBS once more. The cover slips were then placed onto glass slides with mounting media before visualization.

### RNA extraction and quantitative RT-PCR (mammalian cells)

Around 0.6×10^6^ cells were collected. RNA was extracted via the Direct-zol RNA miniprep kit (Zymogen) according to the manufacturer’s instructions. cDNA was generated from 1 mg of total RNA using random hexamers and Maxima Reverse Transcriptase (Thermo Fisher Scientific). Quantitative PCR was performed on the ViiA 7 Real-Time PCR System (Life Technologies) using the Luna qPCR Master Mix (NEB). Thermocycling was done for 40 cycles in a two-step cycling in accordance with the manufacturer’s instructions and each PCR reaction was performed in triplicate. Changes in mRNA expression were quantified using GAPDH and 18s mRNA as references.

### RNA-sequencing

Around 1×10^6^ - 2×10^6^ cells were collected. RNA was extracted via the Direct-zol RNA miniprep kit (Zymogen) according to the manufacturer’s instructions. RNA integrity was confirmed using a 2100 Bioanalyzer (Agilent Technologies). RNA libraries were prepared using the Illumina® Stranded Total RNA Prep, Ligation with Ribo-Zero Plus kit according to manufacturer’s instructions. Illumina NovaSeq 6000 was used by the BU Microarray and Sequencing Core. FastQ files were aligned to the hg38 human genome build using STAR. Ensembl-Gene counts for non-mitochondria genes were generated using featureCounts and Ensembl annotation build 100. The FastQ and alignment quality were assessed using FastQC and RSeQC. Gene expression changes were analyzed using DESeq2. Data were further analyzed with the use of QIAGEN IPA (QIAGEN Inc., https://digitalinsights.qiagen.com/IPA).^147^

### Proteomics Analysis

Proteomics sample preparation and analysis were performed as previously described.^148^ Briefly, protein precipitate was sonicated and reduced using DTT before being digested by sequence-grade trypsin and desalted. Peptides were TMT labelled using the TMT 10-plex isobaric label reagents (Thermofisher) and fractionated on a Waters XBridge BEH C18 (3.5 μm, 4.6 × 250 mm) reverse phase column using an Agilent 1100 HPLC system. Phosphopeptides were enriched using titanium dioxide coated magnetic beads. Fractionated samples and bead-enriched samples were individually loaded onto a C18 trap column (3 μm, 75μm × 2 cm, Thermo Fisher Scientific) connected in-line to a C18 analytical column (2 μm, 75 μm × 50 cm, Thermo EasySpray) using the Thermo EasyLC 1200 system. Peptides were ionized using a nanospray ion source into a Q-Exactive HF mass spectrometer (Thermo Fisher Scientific). MS/MS spectra were searched against the Uniprot human complete proteome FASTA database downloaded on 2018_10_26, using the MaxQuant software (Version 1.6.7.0) that integrates the Andromeda search engine.

### Chromatin immunoprecipitation

For chromatin immunoprecipitation (ChIP) of MDA-MB-231 cells, approximately 2 x 10^7^ cells were cross-linked with 1% formaldehyde at room temperature (∼25°C) for 10 minutes. The cells were then lysed with a hypotonic lysis buffer (10 mM HEPES, 1.5 mM MgCl_2_, 10 mM KCl, 0.5 mM DTT, 0.2 mM PMSF, 1X PIC) for 20 minutes. The nuclei were isolated and lysed with the SDS lysis buffer (1% SDS, 10 mM EDTA, 50 mM Tris-HCl pH 8) for 10 minutes and diluted with ChIP dilution buffer (0.01% SDS, 1.1% Triton-X, 1.2 mM EDTA, 16.7 mM Tris-HCl pH 8, 167 mM NaCl). DNA shearing was performed in Covaris milliTUBE 1 mL with AFA® fiber using the following settings in SonoLab7: Peak Incident Power (PIP) = 140W; Duty Factor = 5%; Cycles per burst (cpb) = 200; time = 480 seconds.

For *C. elegans* ChIP experiments, worms were collected in M9 media and fixed with 2% formaldehyde at 20 °C for 30 minutes. The worms were lysed in RIPA buffer with protease inhibitor cocktail. DNA shearing was performed in Covaris milliTUBE 1 mL with AFA® fiber using the following settings in SonoLab7: Peak Incident Power (PIP) = 240W; Duty Factor = 20%; Cycles per burst (cpb) = 200; time = 480 seconds.

After sonication, 10-25 μg of chromatin was incubated with 1-2 μg of antibodies (Abcam: 9E10, Y69, ab2739, Millipore: 04-835, 05-928, CST: 9402 and anti-MML-1, a generous gift of Dr. Don Ayer, University of Utah) at 4°C overnight. Immunoprecipitated complexes were collected using Protein G Dynabeads (Invitrogen) and washed with 1000 μL of respectively, Low Salt Wash Buffer (150 mM NaCl, 20 mM Tris-HCl pH 8.1, 2 mM EDTA, 1% Triton-X-100, and 0.1% SDS), Medium Salt Wash Buffer (500 mM NaCl, 20 mM Tris-HCl pH 8.1, 2 mM EDTA, 1% Triton-X-100, and 0.1% SDS), High Salt Wash Buffer (1 M NaCl, 20 mM Tris-HCl pH 8.1, 2 mM EDTA, 1% Triton-X-100 and 0.1% SDS), LiCl Wash Buffer (0.25 M LiCl, 10 mM Tris-HCl pH 8.1, 1 mM EDTA, 1% NP40 and 1% sodium deoxycholate), 1X TE (10 mM Tris-HCl pH 8.1, 1mM EDTA). DNA was then extracted and purified by phenol/chloroform extraction. ChIP experiments were repeated at least three times. Significance was calculated by paired Student’s t-test.

The immunoprecipitated DNA was either quantified by qPCR to calculate the percentage of immunoprecipitation relative to the input or subjected to library preparation and high-throughput sequencing.

### MML-1 ChIP-seq Library preparation and analysis

Immunoprecipitated DNA and input libraries (two replicas each) were prepared using TruSeq ChIP Library Preparation kit following the manufacturer’s instructions (Illumina) and checked for concentration and quality on DNA chips with a Bioanalyzer (Agilent). 75-bp single read sequences were generated on the NextSeq 500 sequencer according to manufacturer’s instructions (Illumina).

MML-1 ChIP-seq reads were uploaded to Galaxy and aligned to the C. elegans genome (WS220) using Bowtie (Galaxy Version 1.1.2) with the following parameters: -m 1 --best --strata. Enriched peaks were identified using MACS2 (Galaxy Version 2.1.0.20151222.0) with the following parameters: --gsize 93260000 --bw=175. Statistically significant peaks were filtered using a 5% FDR. Peak coordinates obtained for the two replicates were overlapped, and only overlapping regions were considered. Genomic regions representing ZFP-1 and DOT-1.1 enrichment were fetched from the modENCODE server (http://www.modencode.org). O-GlcNac peaks were obtained from NCBI GEO (GSE18611). For identification of target genes, C. elegans transcription unit coordinates were extracted from the UCSC genome browser (RefSeq based on WS220/ce10). A transcription unit was called bound by MML-1, ZFP-1, DOT-1.1 or O-GlcNac if its 1500bp upstream regions overlapped with a peak. Overlap analysis was performed in R using the package valr.^149^

### In vitro purification

c-MYC – HmuY reporter plasmids and DOT1L C-terminus plasmids were introduced into Rosetta BL21 E. coli according to Heat Shock Transformation protocol and transformed colonies were selected on LB plates containing kanamycin (50 μg/mL). The HmuY DNA sequence was cloned from boiled *P.gingivalis* genomic DNA. A single colony was chosen to grow in 1 – 2 L of LB media containing kanamycin (50 μg/mL). Upon reaching OD600 = 0.4 – 0.8, IPTG was added to the media at a concentration of 400 uM and the culture was incubated at 16 °C overnight. The cultures were pelleted at 4000 x g for 10 minutes, resuspended in the Equilibration Buffer (50 mM Na2PO4, 300 mM NaCl, 10 mM Imidazole) and sonicated three times in a needle sonicator at 70% amplitude for 30 seconds. The expressed protein was then purified using TALON Metal Affinity Resin (Takara Cat. # 635502) according to the manufacturer’s instructions. The purified protein was then buffer exchanged to an imidazole-free phosphate buffer (50 mM Na2PO4, 500 mM NaCl, 5% glycerol) or Tris buffer (50 mM Tris-HCl pH 7.4, 500 mM NaCl, 5% glycerol) and concentrated using the Pierce Protein Concentrator PES, 10 MWCO (ThermoFisher). The resulting purified protein was then aliquoted and stored in −80 °C.

### Recombinant protein cleavage assay

0.5 μg of recombinant human c-Myc (ab169901) was diluted in 20 μL of Buffer G (30 mM tris-HCl pH 7.6, 100 mM NaCl, 10 mM CaCl2, 10 mM MgCl2, 50 μM ATP, 5% glycerol) and incubated with 1 μg of recombinant N-terminal (ab80254) or in vitro purified C-terminal DOT1L protein for 24 hours at 37 °C. The cleaved protein was visualized via western blotting.

For the HmuY reporters, 0.5 μg of in vitro purified HmuY reporter was diluted in 40 μL of Buffer G and incubated with 5 μg of cytoplasmic, nuclear or chromatin lysates or (without protease inhibitor cocktail), or with recombinant N-terminal or in vitro purified C-terminal DOT1L for 24 hours at 37 °C. Cleavage was visualized via western blot.

## Notes

### Competing Interest Statement

The authors have declared no competing interest.

## REFERENCES

1. Poole, C.J., and van Riggelen, J. (2017). MYC—Master Regulator of the Cancer Epigenome and Transcriptome. Genes (Basel) 8, 142. 10.3390/genes8050142.

2. Nair, S.K., and Burley, S.K. (2003). X-Ray Structures of Myc-Max and Mad-Max Recognizing DNA: Molecular Bases of Regulation by Proto-Oncogenic Transcription Factors. Cell 112, 193–205. 10.1016/S0092-8674(02)01284-9.

3. Benassayag, C., Montero, L., Colombié, N., Gallant, P., Cribbs, D., and Morello, D. (2005). Human c-Myc isoforms differentially regulate cell growth and apoptosis in Drosophila melanogaster. Mol Cell Biol 25, 9897–9909. 10.1128/MCB.25.22.9897-9909.2005.

4. Kress, T.R., Sabò, A., and Amati, B. (2015). MYC: connecting selective transcriptional control to global RNA production. Nat Rev Cancer 15, 593–607. 10.1038/nrc3984.

5. Schlosser, I., Hölzel, M., Mürnseer, M., Burtscher, H., Weidle, U.H., and Eick, D. (2003). A role for c-Myc in the regulation of ribosomal RNA processing. Nucleic Acids Res 31, 6148–6156. 10.1093/NAR/GKG794.

6. Tansey, W.P. (2014). Mammalian MYC Proteins and Cancer. New J Sci 2014, 1–27. 10.1155/2014/757534.

7. Carroll, P.A., Freie, B.W., Mathsyaraja, H., and Eisenman, R.N. (2018). The MYC transcription factor network: balancing metabolism, proliferation and oncogenesis. Frontiers of Medicine 2018 12:4 12, 412–425. 10.1007/S11684-018-0650-Z.

8. Billin, A.N., Eilers, A.L., Queva, C., and Ayer, D.E. (1999). Mlx, a Novel Max-like BHLHZip Protein That Interacts with the Max Network of Transcription Factors. Journal of Biological Chemistry 274, 36344–36350. 10.1074/JBC.274.51.36344.

9. Yuan, J., Tirabassi, R.S., Bush, A.B., and Cole, M.D. (1998). The C. elegans MDL-1 and MXL-1 proteins can functionally substitute for vertebrate MAD and MAX. Oncogene 1998 17:9 17, 1109–1118. 10.1038/sj.onc.1202036.

10. Pickett, C.L., Breen, K.T., and Ayer, D.E. (2007). A C. elegans Myc-like network cooperates with semaphorin and Wnt signaling pathways to control cell migration. Dev Biol 310, 226–239. 10.1016/J.YDBIO.2007.07.034.

11. Grove, C.A., De Masi, F., Barrasa, M.I., Newburger, D.E., Alkema, M.J., Bulyk, M.L., and Walhout, A.J.M. (2009). A Multiparameter Network Reveals Extensive Divergence between C. elegans bHLH Transcription Factors. Cell 138, 314–327. 10.1016/J.CELL.2009.04.058.

12. Adhikary, S., and Eilers, M. (2005). Transcriptional regulation and transformation by Myc proteins. Nature Reviews Molecular Cell Biology 2005 6:8 6, 635–645. 10.1038/nrm1703.

13. Cowling, V.H., and Cole, M.D. (2008). An N-Myc truncation analogous to c-Myc-S induces cell proliferation independently of transactivation but dependent on Myc homology box II. Oncogene 27, 1327–1332. 10.1038/sj.onc.1210734.

14. Schuhmacher, M., and Eick, D. (2013). Dose-dependent regulation of target gene expression and cell proliferation by c-Myc levels. Transcription 4, 192–197. 10.4161/TRNS.25907.

15. Xiao, Q., Claassen, G., Shi, J., Adachi, S., Sedivy, J., and Hann, S.R. (1998). Transactivation-defective c-MycS retains the ability to regulate proliferation and apoptosis. Genes Dev 12, 3803–3808.

16. Uribesalgo, I., Buschbeck, M., Gutiérrez, A., Teichmann, S., Demajo, S., Kuebler, B., Nomdedéu, J.F., Marté-N-Caballero, J., Roma, G., Benitah, S.A., et al. (2011). E-box-independent regulation of transcription and differentiation by MYC. Nature Cell Biology 2011 13:12 13, 1443–1449. 10.1038/ncb2355.

17. Ceballos, A., Esse, R., and Grishok, A. (2021). The proline-rich domain of MML-1 is biologically important but not required for localization to target promoters. MicroPubl Biol 2021. 10.17912/MICROPUB.BIOLOGY.000498.

18. Jaenicke, L.A., Rn Von Eyss, B., Carstensen, A., Geyer, M., Eilers, M., and Correspondence, N.P. (2016). Ubiquitin-Dependent Turnover of MYC Antagonizes MYC/PAF1C Complex Accumulation to Drive Transcriptional Elongation. Mol Cell 61, 54–67. 10.1016/j.molcel.2015.11.007.

19. Endres, T., Solvie, D., Heidelberger, J.B., Andrioletti, V., Baluapuri, A., Ade, C.P., Muhar, M., Eilers, U., Vos, S.M., Cramer, P., et al. (2021). Ubiquitylation of MYC couples transcription elongation with double-strand break repair at active promoters. Mol Cell 81, 830–844.e13. 10.1016/J.MOLCEL.2020.12.035.

20. Kim, S.Y., Herbst, A., Tworkowski, K.A., Salghetti, S.E., and Tansey, W.P. (2003). Skp2 Regulates Myc Protein Stability and Activity. Mol Cell 11, 1177–1188. 10.1016/S1097-2765(03)00173-4.

21. Von Der Lehr, N., Johansson, S., Wu, S., Bahram, F., Castell, A., Cetinkaya, C., Hydbring, P., Weidung, I., Nakayama, K., Nakayama, K.I., et al. (2003). The F-Box Protein Skp2 Participates in c-Myc Proteosomal Degradation and Acts as a Cofactor for c-Myc-Regulated Transcription. Mol Cell 11, 1189–1200. 10.1016/S1097-2765(03)00193-X.

22. Alexandrova, E., Salvati, A., Pecoraro, G., Lamberti, J., Melone, V., Sellitto, A., Rizzo, F., Giurato, G., Tarallo, R., Nassa, G., et al. (2022). Histone Methyltransferase DOT1L as a Promising Epigenetic Target for Treatment of Solid Tumors. Front Genet 13, 742. 10.3389/FGENE.2022.864612/BIBTEX.

23. Yang, L., Lei, Q., Li, L., Yang, J., Dong, Z., and Cui, H. (2019). Silencing or inhibition of H3K79 methyltransferase DOT1L induces cell cycle arrest by epigenetically modulating c-Myc expression in colorectal cancer. Clin Epigenetics 11. 10.1186/s13148-019-0778-y.

24. Liu, C., Yang, Q., Zhu, Q., Lu, X., Li, M., Hou, T., Li, Z., Tang, M., Li, Y., Wang, H., et al. (2020). CBP mediated DOT1L acetylation confers DOT1L stability and promotes cancer metastasis. Theranostics 10, 1758–1776. 10.7150/thno.39013.

25. Nassa, G., Salvati, A., Tarallo, R., Gigantino, V., Alexandrova, E., Memoli, D., Sellitto, A., Rizzo, F., Malanga, D., Mirante, T., et al. (2019). Inhibition of histone methyltransferase DOT1L silences ER gene and blocks proliferation of antiestrogen-resistant breast cancer cells. Sci Adv 5. 10.1126/sciadv.aav5590.

26. Vatapalli, R., Sagar, V., Rodriguez, Y., Zhao, J.C., Unno, K., Pamarthy, S., Lysy, B., Anker, J., Han, H., Yoo, Y.A., et al. (2020). Histone methyltransferase DOT1L coordinates AR and MYC stability in prostate cancer. Nat Commun 11, 1–15. 10.1038/s41467-020-18013-7.

27. Zhang, L., Deng, L., Chen, F., Yao, Y., Wu, B., Wei, L., Mo, Q., Song, Y., Zhang, L., Deng, L., et al. (2014). Inhibition of histone H3K79 methylation selectively inhibits proliferation, self-renewal and metastatic potential of breast cancer. Oncotarget 5, 10665–10677. 10.18632/oncotarget.2496.

28. Cho, M.H., Park, J.H., Choi, H.J., Park, M.K., Won, H.Y., Park, Y.J., Lee, C.H., Oh, S.H., Song, Y.S., Kim, H.S., et al. (2015). DOT1L cooperates with the c-Myc-p300 complex to epigenetically derepress CDH1 transcription factors in breast cancer progression. Nat Commun 6. 10.1038/ncomms8821.

29. Liu, D., Zhang, X.X., Li, M.C., Cao, C.H., Wan, D.Y., Xi, B.X., Tan, J.H., Wang, J., Yang, Z.Y., Feng, X.X., et al. (2018). C/EBPβ enhances platinum resistance of ovarian cancer cells by reprogramming H3K79 methylation. Nature Communications 2018 9:1 9, 1–13. 10.1038/s41467-018-03590-5.

30. Chen, C.-W., and Armstrong, S.A. (2015). Targeting DOT1L and HOX gene expression in MLL-rearranged leukemia and beyond. Exp Hematol 43, 673–684. 10.1016/J.EXPHEM.2015.05.012.

31. Zhang, H., Zhou, B., Qin, S., Xu, J., Harding, R., Tempel, W., Nayak, V., Li, Y., Loppnau, P., Dou, Y., et al. (2018). Structural and functional analysis of the DOT1L– AF10 complex reveals mechanistic insights into MLL-AF10-associated leukemogenesis. Genes Dev 32, 341–346. 10.1101/gad.311639.118.

32. Gilan, O., Lam, E.Y.N., Becher, I., Lugo, D., Cannizzaro, E., Joberty, G., Ward, A., Wiese, M., Fong, C.Y., Ftouni, S., et al. (2016). Functional interdependence of BRD4 and DOT1L in MLL leukemia. Nature Structural & Molecular Biology 2016 23:7 23, 673. 10.1038/nsmb.3249.

33. Song, T., Zou, Q., Yan, Y., Lv, S., Li, N., Zhao, X., Ma, X., Liu, H., Tang, B., and Sun, L. (2021). DOT1L O-GlcNAcylation promotes its protein stability and MLL-fusion leukemia cell proliferation. Cell Rep 36, 109739. 10.1016/J.CELREP.2021.109739.

34. Campbell, C.T., Haladyna, J.N., Drubin, D.A., Thomson, T.M., Maria, M.J., Yamauchi, T., Waters, N.J., Olhava, E.J., Pollock, R.M., Smith, J.J., et al. (2017). Mechanisms of pinometostat (EPZ-5676) treatment–emergent resistance in MLL-rearranged leukemia. Mol Cancer Ther 16, 1669–1679. 10.1158/1535-7163.MCT-16-0693.

35. Wong, M., Tee, A.E.L., Milazzo, G., Bell, J.L., Poulos, R.C., Atmadibrata, B., Sun, Y., Jing, D., Ho, N., Ling, D., et al. (2017). The histone methyltransferase DOT1L promotes neuroblastoma by regulating gene transcription. Cancer Res 77, 2522–2533. 10.1158/0008-5472.CAN-16-1663.

36. Guccione, E., Martinato, F., Finocchiaro, G., Luzi, L., Tizzoni, L., Dall’ Olio, V., Zardo, G., Nervi, C., Bernard, L., and Amati, B. (2006). Myc-binding-site recognition in the human genome is determined by chromatin context. Nat Cell Biol 8, 764–770. 10.1038/ncb1434.

37. Steger, D.J., Lefterova, M.I., Ying, L., Stonestrom, A.J., Schupp, M., Zhuo, D., Vakoc, A.L., Kim, J.-E., Chen, J., Lazar, M.A., et al. (2008). DOT1L/KMT4 Recruitment and H3K79 Methylation Are Ubiquitously Coupled with Gene Transcription in Mammalian Cells. Mol Cell Biol 28, 2825–2839. 10.1128/MCB.02076-07.

38. Vlaming, H., and van Leeuwen, F. (2016). The upstreams and downstreams of H3K79 methylation by DOT1L. Chromosoma 2016 125:4 125, 593–605. 10.1007/S00412-015-0570-5.

39. Cao, K., Ugarenko, M., Ozark, P.A., Wang, J., Marshall, S.A., Rendleman, E.J., Liang, K., Wang, L., Zou, L., Smith, E.R., et al. (2020). DOT1L-controlled cell-fate determination and transcription elongation are independent of H3K79 methylation. Proc Natl Acad Sci U S A 117, 27365–27373. 10.1073/pnas.2001075117.

40. Wood, K., Tellier, M., and Murphy, S. (2018). DOT1L and H3K79 Methylation in Transcription and Genomic Stability. Biomolecules 2018, Vol. 8, Page 11 8, 11. 10.3390/BIOM8010011.

41. Nguyen, A.T., and Zhang, Y. (2011). The diverse functions of Dot1 and H3K79 methylation. Genes Dev 25, 1345–1358. 10.1101/GAD.2057811.

42. Cecere, G., Hoersch, S., Jensen, M.B., Dixit, S., and Grishok, A. (2013). The ZFP-1(AF10)/DOT-1 Complex Opposes H2B Ubiquitination to Reduce Pol II Transcription. Mol Cell 50, 894–907. 10.1016/j.molcel.2013.06.002.

43. Wille, C.K., and Sridharan, R. (2022). DOT1L inhibition enhances pluripotency beyond acquisition of epithelial identity and without immediate suppression of the somatic transcriptome. Stem Cell Reports 17, 384–396. 10.1016/J.STEMCR.2021.12.004.

44. Borosha, S., Ratri, A., Ghosh, S., Malcom, C.A., Chakravarthi, V.P., Vivian, J.L., Fields, T.A., Rumi, M.A.K., and Fields, P.E. (2022). DOT1L Mediated Gene Repression in Extensively Self-Renewing Erythroblasts. Front Genet 13, 521. 10.3389/FGENE.2022.828086/BIBTEX.

45. Wu, A., Zhi, J., Tian, T., Cihan, A., Cevher, M.A., Liu, Z., David, Y., Muir, T.W., Roeder, R.G., and Yu, M. (2021). DOT1L complex regulates transcriptional initiation in human erythroleukemic cells. Proc Natl Acad Sci U S A 118, e2106148118. 10.1073/PNAS.2106148118/SUPPL_FILE/PNAS.2106148118.SAPP.PDF.

46. Esse, R., and Grishok, A. (2020). Caenorhabditis elegans Deficient in DOT-1.1 Exhibit Increases in H3K9me2 at Enhancer and Certain RNAi-Regulated Regions. Cells 2020, Vol. 9, Page 1846 9, 1846. 10.3390/CELLS9081846.

47. Avgousti, D.C., Cecere, G., and Grishok, A. (2013). The conserved PHD1-PHD2 domain of ZFP-1/AF10 is a discrete functional module essential for viability in Caenorhabditis elegans. Mol Cell Biol 33, 999–1015. 10.1128/MCB.01462-12.

48. Ran, F.A., Hsu, P.D., Wright, J., Agarwala, V., Scott, D.A., and Zhang, F. (2013). Genome engineering using the CRISPR-Cas9 system. Nat Protoc 8, 2281–2308. 10.1038/nprot.2013.143.

49. Shalem, O., Sanjana, N.E., Hartenian, E., Shi, X., Scott, D.A., Mikkelson, T., Heckl, D., Ebert, B.L., Root, D.E., Doench, J.G., et al. (2014). Genome-scale CRISPR-Cas9 knockout screening in human cells. Science 343, 84–87. 10.1126/science.1247005.

50. Jinek, M., Chylinski, K., Fonfara, I., Hauer, M., Doudna, J.A., and Charpentier, E. (2012). A programmable dual-RNA-guided DNA endonuclease in adaptive bacterial immunity. Science (1979) 337, 816–821. 10.1126/SCIENCE.1225829/SUPPL_FILE/JINEK.SM.PDF.

51. Havula, E., Teesalu, M., Hyötyläinen, T., Seppälä, H., Hasygar, K., Auvinen, P., Orešič, M., Sandmann, T., and Hietakangas, V. (2013). Mondo/ChREBP-Mlx-Regulated Transcriptional Network Is Essential for Dietary Sugar Tolerance in Drosophila. PLoS Genet 9, e1003438. 10.1371/journal.pgen.1003438.

52. O’Donnell, K.A., Wentzel, E.A., Zeller, K.I., Dang, C. v., and Mendell, J.T. (2005). c-Myc-regulated microRNAs modulate E2F1 expression. Nature 2005 435:7043 435, 839–843. 10.1038/nature03677.

53. Aguda, B.D., Kim, Y., Piper-Hunter, M.G., Friedman, A., and Marsh, C.B. (2008). MicroRNA regulation of a cancer network: Consequences of the feedback loops involving miR-17-92, E2F, and Myc. Proc Natl Acad Sci U S A 105, 19678–19683. 10.1073/PNAS.0811166106/ASSET/D0ECBCD1-DDD7-4F25-8B8E-21480AC33915/ASSETS/GRAPHIC/ZPQ9990860800007.JPEG.

54. Li, Y., Choi, P.S., Casey, S.C., Dill, D.L., and Felsher, D.W. (2014). MYC through miR-17-92 suppresses specific target genes to maintain survival, autonomous proliferation, and a Neoplastic state. Cancer Cell 26, 262–272. 10.1016/j.ccr.2014.06.014.

55. Liu, Y.C., Li, F., Handler, J., Huang, C.R.L., Xiang, Y., Neretti, N., Sedivy, J.M., Zeller, K.I., and Dang, C. v. (2008). Global Regulation of Nucleotide Biosynthetic Genes by c-Myc. PLoS One 3, e2722. 10.1371/JOURNAL.PONE.0002722.

56. Muhar, M., Ebert, A., Neumann, T., Umkehrer, C., Jude, J., Wieshofer, C., Rescheneder, P., Lipp, J.J., Herzog, V.A., Reichholf, B., et al. (2018). SLAM-seq defines direct gene-regulatory functions of the BRD4-MYC axis. Science 360, 800–805. 10.1126/science.aao2793.

57. Miyawaki, K., Yamauchi, T., Sugio, T., Sasaki, K., Miyoshi, H., Osborne, S., Taylor, D., Ohshima, K., Kato, K., Maeda, T., et al. (2019). Paics Inhibition Is a Potential Therapeutic Strategy for MYC-Positive DLBCL. Blood 134, 396. 10.1182/BLOOD-2019-129613.

58. Dong, Y., Tu, R., Liu, H., and Qing, G. Regulation of cancer cell metabolism: oncogenic MYC in the driver’s seat. 10.1038/s41392-020-00235-2.

59. Kim, J., Zeller, K.I., Wang, Y., Jegga, A.G., Aronow, B.J., O’Donnell, K.A., and Dang, C. v. (2004). Evaluation of Myc E-Box Phylogenetic Footprints in Glycolytic Genes by Chromatin Immunoprecipitation Assays. Mol Cell Biol 24, 5923–5936. 10.1128/MCB.24.13.5923-5936.2004/SUPPL_FILE/SUPPLEMENTARY_TABLE_S2.DOC.

60. Wang, X., Yu, J., Yan, J., Peng, K., and Zhou, H. (2022). Single-cell sequencing reveals MYC targeting gene MAD2L1 is associated with prostate cancer bone metastasis tumor dormancy. BMC Urol 22, 1–10. 10.1186/S12894-022-00991-Z/FIGURES/8.

61. Menssen, A., Epanchintsev, A., Rezaei, N., Lodygin, D., Jung, P., Verdoodt, B., Diebold, J., and Hermeking, H. (2007). c-MYC delays prometaphase by direct transactivation of MAD2 and BubR1: Identification of mechanisms underlying c-MYC-induced DNA damage and chromosomal instability. Cell Cycle 6, 339–352. 10.4161/CC.6.3.3808.

62. Srikanth, S., Ramachandran, S., and Mohan, S. (2020). Construction of the gene regulatory network identifies MYC as a transcriptional regulator of SWI/SNF complex. Scientific Reports 2020 10:1 10, 1–9. 10.1038/s41598-019-56844-7.

63. Nishimura, K., Fukagawa, T., Takisawa, H., Kakimoto, T., and Kanemaki, M. (2009). An auxin-based degron system for the rapid depletion of proteins in nonplant cells. Nat Methods 6, 917–922. 10.1038/nmeth.1401.

64. Li, S., Prasanna, X., Salo, V.T., Vattulainen, I., and Ikonen, E. (2019). An efficient auxin-inducible degron system with low basal degradation in human cells. Nat Methods 16, 866–869. 10.1038/s41592-019-0512-x.

65. Natsume, T., Kiyomitsu, T., Saga, Y., and Kanemaki, M.T. (2016). Rapid Protein Depletion in Human Cells by Auxin-Inducible Degron Tagging with Short Homology Donors. Cell Rep 15, 210–218. 10.1016/j.celrep.2016.03.001.

66. Helander, S., Montecchio, M., Pilstå, R., and Sears, R.C. (2015). Pre-Anchoring of Pin1 to Unphosphorylated c-Myc in a Fuzzy Complex Regulates c-Myc Activity. Structure 23. 10.1016/j.str.2015.10.010.

67. Farrell, A.S., Pelz, C., Wang, X., Daniel, C.J., Wang, Z., Su, Y., Janghorban, M., Zhang, X., Morgan, C., Impey, S., et al. (2013). Pin1 regulates the dynamics of c-Myc DNA binding to facilitate target gene regulation and oncogenesis. Mol Cell Biol 33, 2930–2949. 10.1128/MCB.01455-12.

68. Chou, T.Y., Hart, G.W., and Dang, C. V (1995). c-Myc is glycosylated at threonine 58, a known phosphorylation site and a mutational hot spot in lymphomas. J Biol Chem 270, 18961–18965. 10.1074/JBC.270.32.18961.

69. Yu, W., Chory, E.J., Wernimont, A.K., Tempel, W., Scopton, A., Federation, A., Marineau, J.J., Qi, J., Barsyte-Lovejoy, D., Yi, J., et al. (2012). Catalytic site remodelling of the DOT1L methyltransferase by selective inhibitors. Nature Communications 2012 3:1 3, 1–12. 10.1038/ncomms2304.

70. Endres, T., Solvie, D., Heidelberger, J.B., Andrioletti, V., Baluapuri, A., Ade, C.P., Muhar, M., Eilers, U., Vos, S.M., Cramer, P., et al. (2021). Ubiquitylation of MYC couples transcription elongation with double-strand break repair at active promoters. Mol Cell 81, 830–844.e13. 10.1016/J.MOLCEL.2020.12.035.

71. JB, H., A, V., ME, B., G, P., S, R., SA, W., and P, B. (2018). Proteomic profiling of VCP substrates links VCP to K6-linked ubiquitylation and c-Myc function. EMBO Rep 19. 10.15252/EMBR.201744754.

72. de Almeida, M., Hinterndorfer, M., Brunner, H., Grishkovskaya, I., Singh, K., Schleiffer, A., Jude, J., Deswal, S., Kalis, R., Vunjak, M., et al. (2021). AKIRIN2 controls the nuclear import of proteasomes in vertebrates. Nature 2021 599:7885 599, 491–496. 10.1038/s41586-021-04035-8.

73. Geng, F., Wenzel, S., and Tansey, W.P. (2012). Ubiquitin and Proteasomes in Transcription. https://doi.org/10.1146/annurev-biochem-052110-120012 81, 177–201. 10.1146/ANNUREV-BIOCHEM-052110-120012.

74. Johnson, D.W., Llop, J.R., Farrell, S.F., Yuan, J., Stolzenburg, L.R., and Samuelson, A. V. (2014). The Caenorhabditis elegans Myc-Mondo/Mad Complexes Integrate Diverse Longevity Signals. PLoS Genet 10, e1004278. 10.1371/JOURNAL.PGEN.1004278.

75. Shimada, M., Kanematsu, K., Tanaka, K., Yokosawa, H., and Kawahara, H. (2006). Proteasomal ubiquitin receptor RPN-10 controls sex determination in Caenorhabditis elegans. Mol Biol Cell 17, 5356–5371. 10.1091/MBC.E06-05-0437/ASSET/IMAGES/LARGE/ZMK0120678950012.JPEG.

76. Keith, S.A., Maddux, S.K., Zhong, Y., Chinchankar, M.N., Ferguson, A.A., Ghazi, A., and Fisher, A.L. (2016). Graded Proteasome Dysfunction in Caenorhabditis elegans Activates an Adaptive Response Involving the Conserved SKN-1 and ELT-2 Transcription Factors and the Autophagy-Lysosome Pathway. PLoS Genet 12, e1005823. 10.1371/JOURNAL.PGEN.1005823.

77. Hyer, M.L., Milhollen, M.A., Ciavarri, J., Fleming, P., Traore, T., Sappal, D., Huck, J., Shi, J., Gavin, J., Brownell, J., et al. (2018). A small-molecule inhibitor of the ubiquitin activating enzyme for cancer treatment. Nat Med 24, 186–193. 10.1038/nm.4474.

78. Solvie, D., Baluapuri, A., Uhl, L., Fleischhauer, D., Endres, T., Papadopoulos, D., Aziba, A., Gaballa, A., Mikicic, I., Isaakova, E., et al. (2022). MYC multimers shield stalled replication forks from RNA polymerase. Nature 2022 612:7938 612, 148–155. 10.1038/s41586-022-05469-4.

79. Meyer, H., and van den Boom, J. (2023). Targeting of client proteins to the VCP/p97/Cdc48 unfolding machine. Front Mol Biosci 10, 76. 10.3389/FMOLB.2023.1142989/BIBTEX.

80. Meyer, H., Bug, M., and Bremer, S. (2012). Emerging functions of the VCP/p97 AAA-ATPase in the ubiquitin system. Nature Cell Biology 2012 14:2 14, 117–123. 10.1038/ncb2407.

81. Radhakrishnan, S.K., den Besten, W., and Deshaies, R.J. (2014). p97-dependent retrotranslocation and proteolytic processing govern formation of active Nrf1 upon proteasome inhibition. Elife 2014. 10.7554/ELIFE.01856.001.

82. Krastev, D.B., Li, S., Sun, Y., Wicks, A.J., Hoslett, G., Weekes, D., Badder, L.M., Knight, E.G., Marlow, R., Pardo, M.C., et al. (2022). The ubiquitin-dependent ATPase p97 removes cytotoxic trapped PARP1 from chromatin. Nature Cell Biology 2022 24:1 24, 62–73. 10.1038/s41556-021-00807-6.

83. Torrecilla, I., Oehler, J., and Ramadan, K. (2017). The role of ubiquitin-dependent segregase p97 (VCP or Cdc48) in chromatin dynamics after DNA double strand breaks. Philosophical Transactions of the Royal Society B: Biological Sciences 372. 10.1098/RSTB.2016.0282.

84. Lehrbach, N.J., and Ruvkun, G. (2016). Proteasome dysfunction triggers activation of SKN-1A/Nrf1 by the aspartic protease DDI-1. Elife 5. 10.7554/ELIFE.17721.001.

85. Radhakrishnan, S.K., den Besten, W., and Deshaies, R.J. (2014). p97-dependent retrotranslocation and proteolytic processing govern formation of active Nrf1 upon proteasome inhibition. Elife 2014. 10.7554/ELIFE.01856.001.

86. Wang, X., Sato, R., Brown, M.S., Hua, X., and Goldstein, J.L. (1994). SREBP-1, a membrane-bound transcription factor released by sterol-regulated proteolysis. Cell 77, 53–62. 10.1016/0092-8674(94)90234-8.

87. Varshavsky, A. (2011). The N-end rule pathway and regulation by proteolysis. Protein Sci 20, 1298. 10.1002/PRO.666.

88. Piatkov, K.I., Brower, C.S., and Varshavsky, A. (2012). The N-end rule pathway counteracts cell death by destroying proapoptotic protein fragments. Proc Natl Acad Sci U S A 109, E1839–E1847. 10.1073/PNAS.1207786109/SUPPL_FILE/PNAS.201207786SI.PDF.

89. Xu, Z., Payoe, R., and Fahlman, R.P. (2012). The C-terminal Proteolytic Fragment of the Breast Cancer Susceptibility Type 1 Protein (BRCA1) Is Degraded by the N-end Rule Pathway. Journal of Biological Chemistry 287, 7495–7502. 10.1074/JBC.M111.301002.

90. Heo, A.J., Kim, S., Bin, Kwon, Y.T., and Ji, C.H. (2023). The N-degron pathway: From basic science to therapeutic applications. Biochimica et Biophysica Acta (BBA) - Gene Regulatory Mechanisms 1866, 194934. 10.1016/J.BBAGRM.2023.194934.

91. Koizumi, S., Irie, T., Hirayama, S., Sakurai, Y., Yashiroda, H., Naguro, I., Ichijo, H., Hamazaki, J., and Murata, S. (2016). The aspartyl protease DDI2 activates Nrf1 to compensate for proteasome dysfunction. Elife 5. 10.7554/ELIFE.18357.

92. Dirac-Svejstrup, A.B., Walker, J., Faull, P., Encheva, V., Akimov, V., Puglia, M., Perkins, D., Kümper, S., Hunjan, S.S., Blagoev, B., et al. (2020). DDI2 Is a Ubiquitin-Directed Endoprotease Responsible for Cleavage of Transcription Factor NRF1. Mol Cell 79, 332. 10.1016/J.MOLCEL.2020.05.035.

93. Yip, M.C.J., Bodnar, N.O., and Rapoport, T.A. (2020). Ddi1 is a ubiquitin-dependent protease. Proc Natl Acad Sci U S A 117, 7776–7781. 10.1073/PNAS.1902298117/-/DCSUPPLEMENTAL.

94. Kagaya, S., Kitanaka, C., Noguchi, K., Mochizuki, T., Sugiyama, A., Asai, A., Yasuhara, N., Eguchi, Y., Tsujimoto, Y., and Kuchino, Y. (1997). A Functional Role for Death Proteases in s-Myc-and c-Myc-Mediated Apoptosis.

95. Chen, L., Smith, L., Accavitti-Loper, M.A., Omura, S., and Bingham Smith, J. (2000). Ubiquitylation and Destruction of Endogenous c-MycS by the Proteasome: Are Myc Boxes Dispensable? Arch Biochem Biophys 374, 306–312. 10.1006/ABBI.1999.1603.

96. Olczak, T., Sroka, A., Potempa, J., and Olczak, M. (2008). Porphyromonas gingivalis HmuY and HmuR: Further characterization of a novel mechanism of heme utilization. Arch Microbiol 189, 197–210. 10.1007/s00203-007-0309-7.

97. Wójtowicz, H., Guevara, T., Tallant, C., Olczak, M., Sroka, A., Potempa, J., Solà, M., Olczak, T., and Gomis-Rüth, F.X. (2009). Unique Structure and Stability of HmuY, a Novel Heme-Binding Protein of Porphyromonas gingivalis. PLoS Pathog 5, e1000419. 10.1371/journal.ppat.1000419.

98. Falkowski, K., Bielecka, E., Thøgersen, I.B., Bocheńska, O., Płaza, K., Kalińska, M., Sąsiadek, L., Magoch, M., Pęcak, A., Wiśniewska, M., et al. (2020). Kallikrein-Related Peptidase 14 Activates Zymogens of Membrane Type Matrix Metalloproteinases (MT-MMPs)—A CleavEx Based Analysis. International Journal of Molecular Sciences 2020, Vol. 21, Page 4383 21, 4383. 10.3390/IJMS21124383.

99. Conacci-Sorrell, M., Ngouenet, C., and Eisenman, R.N. (2010). Myc-nick: A cytoplasmic cleavage product of Myc that promotes α-tubulin acetylation and cell differentiation. Cell 142, 480–493. 10.1016/j.cell.2010.06.037.

100. Conacci-Sorrell, M., Ngouenet, C., Anderson, S., Brabletz, T., and Eisenman, R.N. (2014). Stress-induced cleavage of Myc promotes cancer cell survival. Genes Dev 28, 689–707. 10.1101/GAD.231894.113.

101. Navia, M.A., Fitzgerald, P.M.D., McKeever, B.M., Leu, C.T., Heimbach, J.C., Herber, W.K., Sigal, I.S., Darke, P.L., and Springer, J.P. (1989). Three-dimensional structure of aspartyl protease from human immunodeficiency virus HIV-1. Nature 1989 337:6208 337, 615–620. 10.1038/337615a0.

102. Min, J., Feng, Q., Li, Z., Zhang, Y., and Xu, R.M. (2003). Structure of the catalytic domain of human Dot1L, a non-SET domain nucleosomal histone methyltransferase. Cell 112, 711–723. 10.1016/S0092-8674(03)00114-4.

103. Capotosti, F., Guernier, S., Lammers, F., Waridel, P., Cai, Y., Jin, J., Conaway, J.W., Conaway, R.C., and Herr, W. (2011). O-GlcNAc transferase catalyzes site-specific proteolysis of HCF-1. Cell 144, 376–388. 10.1016/j.cell.2010.12.030.

104. Lazarus, M.B., Jiang, J., Kapuria, V., Bhuiyan, T., Janetzko, J., Zandberg, W.F., Vocadlo, D.J., Herr, W., and Walker, S. (2013). HCF-1 is cleaved in the active site of O-GlcNAc transferase. Science 342, 1235–1239. 10.1126/science.1243990.

105. Kim, S.K., Jung, I., Lee, H., Kang, K., Kim, M., Jeong, K., Kwon, C.S., Han, Y.M., Kim, Y.S., Kim, D., et al. (2012). Human histone H3K79 methyltransferase DOT1L methyltransferase binds actively transcribing RNA polymerase II to regulate gene expression. Journal of Biological Chemistry 287, 39698–39709. 10.1074/jbc.M112.384057.

106. van Welsem, T., Korthout, T., Ekkebus, R., Morais, D., Molenaar, T.M., van Harten, K., Poramba-Liyanage, D.W., Sun, S.M., Lenstra, T.L., Srivas, R., et al. (2018). Dot1 promotes H2B ubiquitination by a methyltransferase-independent mechanism. Nucleic Acids Res 46, 11251–11261. 10.1093/NAR/GKY801.

107. Deshpande, A.J., Deshpande, A., Sinha, A.U., Chen, L., Chang, J., Cihan, A., Fazio, M., Chen, C.-W., Zhu, N., Koche, R., et al. (2014). Article AF10 Regulates Progressive H3K79 Methylation and HOX Gene Expression in Diverse AML Subtypes. 10.1016/j.ccell.2014.10.009.

108. Chen, C.W., Koche, R.P., Sinha, A.U., Deshpande, A.J., Zhu, N., Eng, R., Doench, J.G., Xu, H., Chu, S.H., Qi, J., et al. (2015). DOT1L inhibits SIRT1-mediated epigenetic silencing to maintain leukemic gene expression in MLL-rearranged leukemia. Nature Medicine 2015 21:4 21, 335–343. 10.1038/nm.3832.

109. Waters, N.J. (2017). Preclinical Pharmacokinetics and Pharmacodynamics of Pinometostat (EPZ-5676), a First-in-Class, Small Molecule S-Adenosyl Methionine Competitive Inhibitor of DOT1L. Eur J Drug Metab Pharmacokinet 42, 891–901. 10.1007/S13318-017-0404-3/TABLES/3.

110. Nassa, G., Salvati, A., Tarallo, R., Gigantino, V., Alexandrova, E., Memoli, D., Sellitto, A., Rizzo, F., Malanga, D., Mirante, T., et al. (2019). Inhibition of histone methyltransferase DOT1L silences ERα gene and blocks proliferation of antiestrogen-resistant breast cancer cells. Sci Adv 5, eaav5590. 10.1126/sciadv.aav5590.

111. Oktyabri, D., Ishimura, A., Tange, S., Terashima, M., and Suzuki, T. (2016). DOT1L histone methyltransferase regulates the expression of BCAT1 and is involved in sphere formation and cell migration of breast cancer cell lines. Biochimie 123, 20–31. 10.1016/J.BIOCHI.2016.01.005.

112. Salvati, A., Gigantino, V., Nassa, G., Giurato, G., Alexandrova, E., Rizzo, F., Tarallo, R., and Weisz, A. (2019). The Histone Methyltransferase DOT1L Is a Functional Component of Estrogen Receptor Alpha Signaling in Ovarian Cancer Cells. Cancers 2019, Vol. 11, Page 1720 11, 1720. 10.3390/CANCERS11111720.

113. Zhang, X., Liu, D., Li, M., Cao, C., Wan, D., Xi, B., Li, W., Tan, J., Wang, J., Wu, Z., et al. (2017). Prognostic and therapeutic value of disruptor of telomeric silencing-1-like (DOT1L) expression in patients with ovarian cancer. J Hematol Oncol 10, 1–13. 10.1186/S13045-017-0400-8/FIGURES/6.

114. Alexandrova, E., Lamberti, J., Memoli, D., Quercia, C., Melone, V., Rizzo, F., Tarallo, R., Giurato, G., Nassa, G., and Weisz, A. (2022). Combinatorial targeting of menin and the histone methyltransferase DOT1L as a novel therapeutic strategy for treatment of chemotherapy-resistant ovarian cancer. Cancer Cell Int 22, 1–13. 10.1186/S12935-022-02740-6/FIGURES/4.

115. Kryczek, I., Lin, Y., Nagarsheth, N., Peng, D., Zhao, L., Zhao, E., Vatan, L., Szeliga, W., Dou, Y., Owens, S., et al. (2014). IL-22+CD4+ T Cells Promote Colorectal Cancer Stemness via STAT3 Transcription Factor Activation and Induction of the Methyltransferase DOT1L. Immunity 40, 772–784. 10.1016/J.IMMUNI.2014.03.010.

116. Campbell, J.D., Alexandrov, A., Kim, J., Wala, J., Berger, A.H., Pedamallu, C.S., Shukla, S.A., Guo, G., Brooks, A.N., Murray, B.A., et al. (2016). Distinct patterns of somatic genome alterations in lung adenocarcinomas and squamous cell carcinomas. Nature Genetics 2016 48:6 48, 607–616. 10.1038/ng.3564.

117. Loeser, H., Waldschmidt, D., Kuetting, F., Heydt, C., Zander, T., Plum, P., Alakus, H., Buettner, R., and Quaas, A. (2017). Copy-number variation and protein expression of DOT1L in pancreatic adenocarcinoma as a potential drug target. Mol Clin Oncol 6, 639–642. 10.3892/MCO.2017.1194.

118. von Mikecz, A. (2006). The nuclear ubiquitin-proteasome system. J Cell Sci 119, 1977–1984. 10.1242/JCS.03008.

119. Song, C., Wang, Q., Song, C., Lockett, S.J., Colburn, N.H., Li, C.C.H., Wang, J.M., and Rogers, T.J. (2015). Nucleocytoplasmic Shuttling of Valosin-Containing Protein (VCP/p97) Regulated by Its N domain and C-terminal Region. Biochim Biophys Acta 1853, 222. 10.1016/J.BBAMCR.2014.10.019.

120. Lipford, J.R., Smith, G.T., Chi, Y., and Deshaies, R.J. (2005). A putative stimulatory role for activator turnover in gene expression. Nature 2005 438:7064 438, 113–116. 10.1038/nature04098.

121. Molinari, E., Gilman, M., and Natesan, S. (1999). Proteasome-mediated degradation of transcriptional activators correlates with activation domain potency in vivo. EMBO J 18, 6439–6447. 10.1093/EMBOJ/18.22.6439.

122. Salghetti, S.E., Muratani, M., Wijnen, H., Futcher, B., and Tansey, W.P. (2000). Functional overlap of sequences that activate transcription and signal ubiquitin-mediated proteolysis. Proc Natl Acad Sci U S A 97, 3118–3123. 10.1073/PNAS.97.7.3118/ASSET/12C88155-DB08-4198-BBE9-9A4D41773A4D/ASSETS/GRAPHIC/PQ0500075005.JPEG.

123. Sundqvist, A., and Ericsson, J. (2003). Transcription-dependent degradation controls the stability of the SREBP family of transcription factors. Proc Natl Acad Sci U S A 100, 13833–13838. 10.1073/PNAS.2335135100/SUPPL_FILE/5135FIG10.PDF.

124. Nie, Z., Guo, C., Das, S.K., Chow, C.C., Batchelor, E., Simons Jnr, S.S., and Levens, D. (2020). Dissecting transcriptional amplification by MYC. Elife 9, 1–32. 10.7554/eLife.52483.

125. Ahlstedt, B.A., Ganji, R., and Raman, M. (2022). The functional importance of VCP to maintaining cellular protein homeostasis. Biochem Soc Trans 50, 1457. 10.1042/BST20220648.

126. Franz, A., Ackermann, L., and Hoppe, T. (2016). Ring of change: CDC48/p97 drives protein dynamics at chromatin. Front Genet 7, 194837. 10.3389/FGENE.2016.00073/BIBTEX.

127. Dantuma, N.P., and Hoppe, T. (2012). Growing sphere of influence: Cdc48/p97 orchestrates ubiquitin-dependent extraction from chromatin. Trends Cell Biol 22, 483–491. 10.1016/j.tcb.2012.06.003.

128. Kröning, A., van den Boom, J., Kracht, M., Kueck, A.F., and Meyer, H. (2022). Ubiquitin-directed AAA+ ATPase p97/VCP unfolds stable proteins crosslinked to DNA for proteolysis by SPRTN. Journal of Biological Chemistry 298, 101976. 10.1016/J.JBC.2022.101976.

129. Lehner, M.H., Walker, J., Temcinaite, K., Herlihy, A., Taschner, M., Berger, A.C., Corbett, A.H., Dirac Svejstrup, A.B., and Svejstrup, J.Q. (2022). Yeast Smy2 and its human homologs GIGYF1 and -2 regulate Cdc48/VCP function during transcription stress. Cell Rep 41, 111536. 10.1016/j.celrep.2022.111536.

130. Verma, R., Oania, R., Fang, R., Smith, G.T., and Deshaies, R.J. (2011). Cdc48/p97 mediates UV-dependent turnover of RNA Pol II. Mol Cell 41, 82–92. 10.1016/j.molcel.2010.12.017.

131. Wilcox, A.J., and Laney, J.D. (2009). A ubiquitin-selective AAA-ATPase mediates transcriptional switching by remodelling a repressor–promoter DNA complex. Nature Cell Biology 2009 11:12 11, 1481–1486. 10.1038/ncb1997.

132. Alexandru, G., Graumann, J., Smith, G.T., Kolawa, N.J., Fang, R., and Deshaies, R.J. (2008). UBXD7 binds multiple ubiquitin ligases and implicates p97 in HIF1α turnover. Cell 134, 804–816. 10.1016/j.cell.2008.06.048.

133. Anderson, S., Poudel, K.R., Roh-Johnson, M., Brabletz, T., Yu, M., Borenstein-Auerbach, N., Grady, W.N., Bai, J., Moens, C.B., Eisenman, R.N., et al. (2016). MYC-nick promotes cell migration by inducing fascin expression and Cdc42 activation. Proc Natl Acad Sci U S A 113, E5481–E5490. 10.1073/PNAS.1610994113/SUPPL_FILE/PNAS.201610994SI.PDF.

134. Shoji, W., Suenaga, Y., Kaneko, Y., Islam, S.M.R., Alagu, J., Yokoi, S., Nio, M., and Nakagawara, A. (2015). NCYM promotes calpain-mediated Myc-nick production in human MYCN-amplified neuroblastoma cells. Biochem Biophys Res Commun 461, 501–506. 10.1016/J.BBRC.2015.04.050.

135. Allen-Petersen, B.L., and Sears, R.C. (2019). Mission Possible: Advances in MYC Therapeutic Targeting in Cancer. Preprint at Springer International Publishing, 10.1007/s40259-019-00370-5 10.1007/s40259-019-00370-5.

136. Beaulieu, M.E., Jauset, T., Massó-Vallés, D., Martínez-Martín, S., Rahl, P., Maltais, L., Zacarias-Fluck, M.F., Casacuberta-Serra, S., del Pozo, E.S., Fiore, C., et al. (2019). Intrinsic cell-penetrating activity propels omomyc from proof of concept to viable anti-myc therapy. Sci Transl Med 11. 10.1126/scitranslmed.aar5012.

137. Struntz, N.B., Chen, A., Deutzmann, A., Wilson, R.M., Stefan, E., Evans, H.L., Ramirez, M.A., Liang, T., Caballero, F., Wildschut, M.H.E., et al. (2019). Stabilization of the Max Homodimer with a Small Molecule Attenuates Myc-Driven Transcription. Cell Chem Biol 26, 711–723.e14. 10.1016/j.chembiol.2019.02.009.

138. Whitfield, J.R., Beaulieu, M.E., and Soucek, L. (2017). Strategies to inhibit Myc and their clinical applicability. Preprint at Frontiers Media S.A., 10.3389/fcell.2017.00010 10.3389/fcell.2017.00010.

139. Brenner, S. (1974). THE GENETICS OF CAENORHABDITIS ELEGANS. Genetics 77, 71–94. 10.1093/GENETICS/77.1.71.

140. Irizarry, R. (2012). Selected Works of Terry Speed.

141. Benjamini, Y., and Hochberg, Y. (1995). Controlling the False Discovery Rate: A Practical and Powerful Approach to Multiple Testing. Journal of the Royal Statistical Society: Series B (Methodological) 57, 289–300. 10.1111/j.2517-6161.1995.tb02031.x.

142. Tresse, E., Salomons, F.A., Vesa, J., Bott, L.C., Kimonis, V., Yao, T.P., Dantuma, N.P., and Taylor, J.P. (2010). VCP/p97 is essential for maturation of ubiquitin-containing autophagosomes and this function is impaired by mutations that cause IBMPFD. Autophagy 6, 217–227. 10.4161/AUTO.6.2.11014.

143. Sanjana, N.E., Shalem, O., and Zhang, F. (2014). Improved vectors and genome-wide libraries for CRISPR screening. Nat Methods 11, 783–784. 10.1038/nmeth.3047.

144. Cong, L., Ran, F.A., Cox, D., Lin, S., Barretto, R., Habib, N., Hsu, P.D., Wu, X., Jiang, W., Marraffini, L.A., et al. (2013). Multiplex Genome Engineering Using CRISPR/Cas Systems. Science (1979) 339, 819–823. 10.1126/science.1231143.

145. Sarbassov, D.D., Guertin, D.A., Ali, S.M., and Sabatini, D.M. (2005). Phosphorylation and regulation of Akt/PKB by the rictor-mTOR complex. Science 307, 1098–1101. 10.1126/science.1106148.

146. Gillotin, S. (2018). Isolation of Chromatin-bound Proteins from Subcellular Fractions for Biochemical Analysis. Bio Protoc 8. 10.21769/bioprotoc.3035.

147. Krämer, A., Green, J., Pollard, J., and Tugendreich, S. (2014). Causal analysis approaches in Ingenuity Pathway Analysis. Bioinformatics 30, 523–530. 10.1093/bioinformatics/btt703.

148. Chan, S., Smith, E., Gao, Y., Kwan, J., Blum, B.C., Tilston-Lunel, A.M., Turcinovic, I., Varelas, X., Cardamone, M.D., Monti, S., et al. (2020). Loss of G-Protein Pathway Suppressor 2 Promotes Tumor Growth Through Activation of AKT Signaling. Front Cell Dev Biol 8, 608044. 10.3389/fcell.2020.608044.

149. Riemondy, K.A., Sheridan, R.M., Gillen, A., Yu, Y., Bennett, C.G., and Hesselberth, J.R. (2017). valr: Reproducible genome interval analysis in R. F1000Res 6, 1025. 10.12688/f1000research.11997.1.

